# RTF2 controls replication repriming and ribonucleotide excision at the replisome

**DOI:** 10.1101/2023.03.13.532415

**Authors:** Brooke A. Conti, Penelope D. Ruiz, Cayla Broton, Nicolas J. Blobel, Molly C. Kottemann, Sunandini Sridhar, Francis P. Lach, Tom Wiley, Nanda K. Sasi, Thomas Carroll, Agata Smogorzewska

## Abstract

Genetic information is duplicated via the highly regulated process of DNA replication. The machinery coordinating this process, the replisome, encounters many challenges, including replication fork-stalling lesions that threaten the accurate and timely transmission of genetic information. Cells have multiple mechanisms to repair or bypass lesions that would otherwise compromise DNA replication^1,2^. We have previously shown that proteasome shuttle proteins, DNA Damage Inducible 1 and 2 (DDI1/2) function to regulate Replication Termination Factor 2 (RTF2) at the stalled replisome, allowing for replication fork stabilization and restart^3^. Here we show that RTF2 regulates replisome localization of RNase H2, a heterotrimeric enzyme responsible for removing RNA in the context of RNA-DNA heteroduplexes^4–6^. We show that during unperturbed DNA replication, RTF2, like RNase H2, is required to maintain normal replication fork speeds. However, persistent RTF2 and RNase H2 at stalled replication forks compromises the replication stress response, preventing efficient replication restart. Such restart is dependent on PRIM1, the primase component of DNA polymerase α-primase. Our data show a fundamental need for regulation of replication-coupled ribonucleotide incorporation during normal replication and the replication stress response that is achieved through RTF2. We also provide evidence for PRIM1 function in direct replication restart following replication stress in mammalian cells.

## Main text

### RTF2 is necessary for *in vivo* viability, cellular proliferation, and DNA replication

To investigate the function of RTF2, we intercrossed *Rtf2^+/-^*and *Rtf2^+/stop^* mice (Extended Data Fig. 1a-c). While heterozygous mice were viable and grew at rates comparable to those of wild type mice, neither *Rtf2^stop/stop^* nor *Rtf2^-/-^* offspring were observed, indicating that RTF2 is essential for embryonic development (Extended Data Fig. 1d-f). *Rtf2^-/lox^* mouse embryonic fibroblasts (MEFs) treated with Cre recombinase replicated slowly, a phenotype that was rescued by complementation with expression of mRTF2 (Fig. 1a,b, Extended Data Fig. 2a-g). Based on the *in vivo* and *in vitro* data, we conclude that RTF2 is required for organismal and cellular growth.

**Figure 1.**
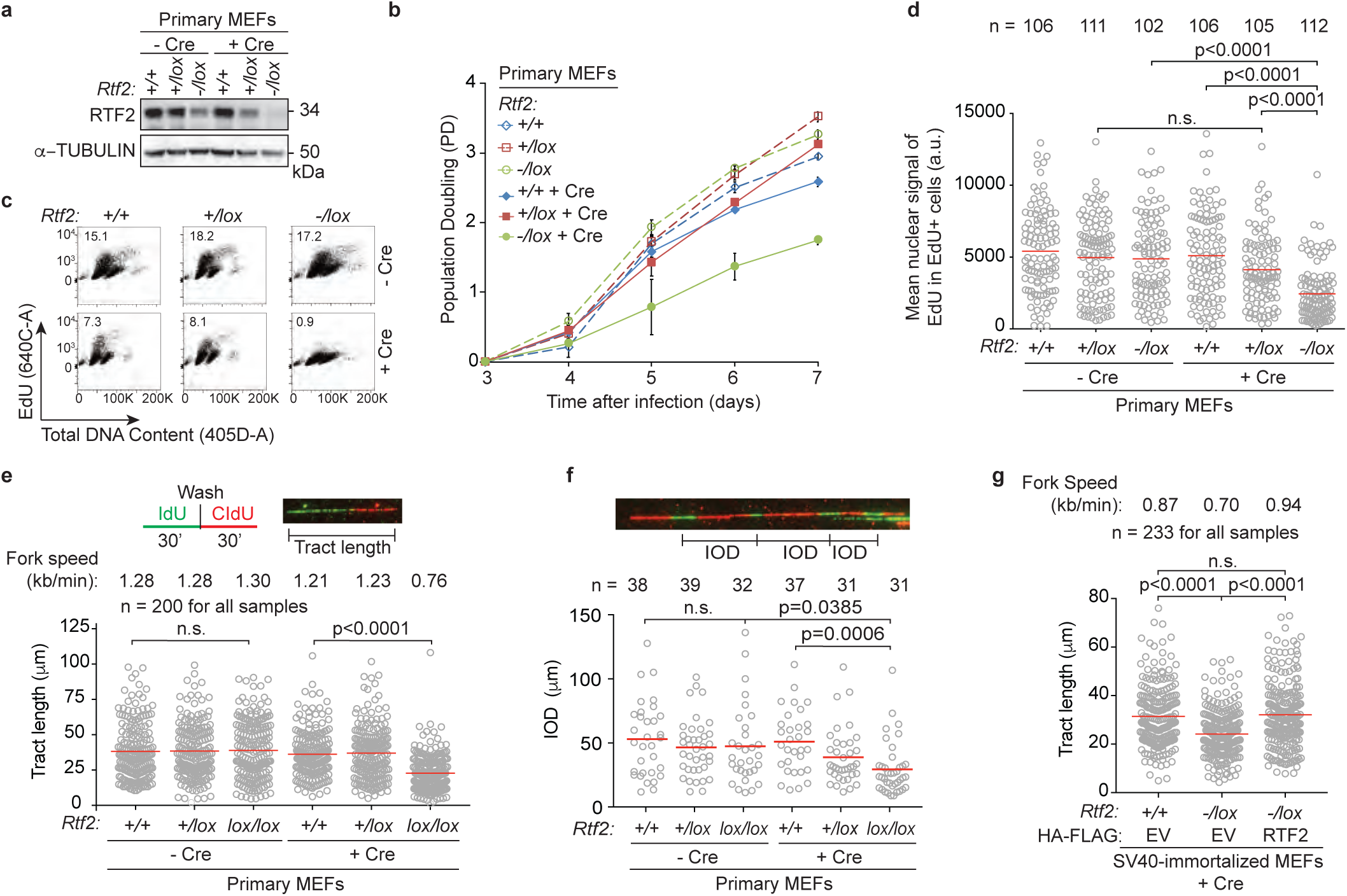
RTF2 is necessary for cellular proliferation and DNA replication. **a**, Representative immunoblot of whole cell lysates showing RTF2 levels 72 hrs after transduction of MEFs with Hit & Run pMMP Cre retrovirus^41^. **b,** Representative growth curves in MEFs transduced with pWZL Cre-hygro retrovirus. **c,** Representative cell cycle profiles from flow cytometry at 72 hrs after Cre. **d,** Quantification of representative experiment of mean nuclear signal of EdU from EdU-positive cells at 72 hrs after Cre. **e,** Top: Schematic and representative image of DNA combing. Below: Quantification of representative experiment of replication tract lengths from primary MEFs at 72 hrs after Cre. **f,** Top: Representative image of inter-origin distances measured within replication clusters. Bottom: Quantification of representative experiment of inter-origin distances (IOD) from primary MEFs at 72 hrs after Cre. **g,** Quantification of representative experiment of replication tract lengths from SV40-immortalized MEFs stably expressing HA-FLAG empty vector (EV) or Rtf2 at 120 hrs after Cre. Error bars represent standard deviation for b. Experiments were conducted at least three times in biological replicates with consistent results for a-g. Mean shown with red line for d,e,f,g. Experiments were blinded prior to analysis for d,e,f,g. Average fork speeds are listed above each sample for e and g. Outliers removed with ROUT (1%) for e and g. Significance was evaluated by Kruskal-Wallis ANOVA with a Dunn’s post-test.

RTF2-deficient cells exhibited abnormal nuclear morphology, including multinucleated cells and cells with multiple micronuclei (Extended Data Fig. 2h,i). Live cell imaging of RTF2-deficient cells expressing GFP-H2B revealed an increased percentage of abnormal mitoses in the absence of RTF2 (Extended Data Fig. 2j,k Supplementary Movies 1-3). The accumulation of abnormal nuclei was suppressed by treatment with RO-3306, a potent inhibitor of CDK1^7^ that arrests cells in G2 and prevents entry into M phase (Extended Data Fig. 2l,m), underscoring that the increase in abnormal nuclei is the result of aberrant cellular division.

Given that RTF2 is a component of the replisome, we hypothesized that aberrant cellular division in the setting of RTF2 loss could follow aberrant DNA replication^3,8,9^. We assessed cell cycle profiles of cells lacking RTF2 with 5-Ethynyl-2-deoxyuridine (EdU) labeling (Fig. 1c, Extended Data Fig. 3a,b). RTF2-deficient cells displayed a decrease in the percentage of S phase cells, and cells in S phase had lower peak EdU intensity (Fig. 1c,d, Extended Data Fig. 3c-f). DNA combing revealed a significant decrease in replication speed and shortening of the inter-origin distance in RTF2-deficient cells (Fig. 1e,f). The replication speed defect was complemented by expression of mRTF2 cDNA (Fig. 1g, Extended Data Fig. 3g). The replication tracks emanating bidirectionally from an origin of replication were symmetrical (Extended Data Fig. 3h), suggesting that, though slow-moving, RTF2-deficient replication forks are stable.

Previous reports in *Arabidopsis thaliana* suggest RTF2 has a role in intron retention, likely through a non-conserved N-terminal extension in the atRTF2 protein. We did not detect any significant changes in global intron retention, and the transcriptional profiles of RTF2-deficient MEFs were largely unchanged (Extended Data Fig. 4a-c)^10^.

### RTF2 function during unperturbed replication is dependent on RNase H2

To identify the mechanism responsible for the observed replication defect, we employed isolation of proteins on nascent DNA (iPOND) to examine the changes in the replication fork proteome in the absence of RTF2 (Fig. 2a)^11^. While many replisome components were unchanged, peptides from the RNase H2 complex were lost from replication forks in RTF2-deficient MEFs (Fig. 2b,c). Nascent DNA proximity ligation assay (nPLA), an orthogonal method to detect the presence of proteins at nascent DNA^12^, showed a dramatic decrease of endogenous RNASEH2A, the catalytic subunit of RNase H2, at the replication fork in the absence of RTF2 (Fig. 2d, Extended Data Fig. 5a). The decrease in RNASEH2A at the replication fork is not due to loss of *Rnaseh2a* transcripts or total protein levels (Extended Data Fig. 5b-d). Increasing the levels of RTF2 at replication forks through DDI1/2 depletion led to increased levels of RNase H2 at nascent chromatin (Extended Data Fig. 5e)^3^. RNase H2 loss, however, did not result in loss of RTF2 from replication forks (Extended Data Fig. 5f).

**Figure 2.**
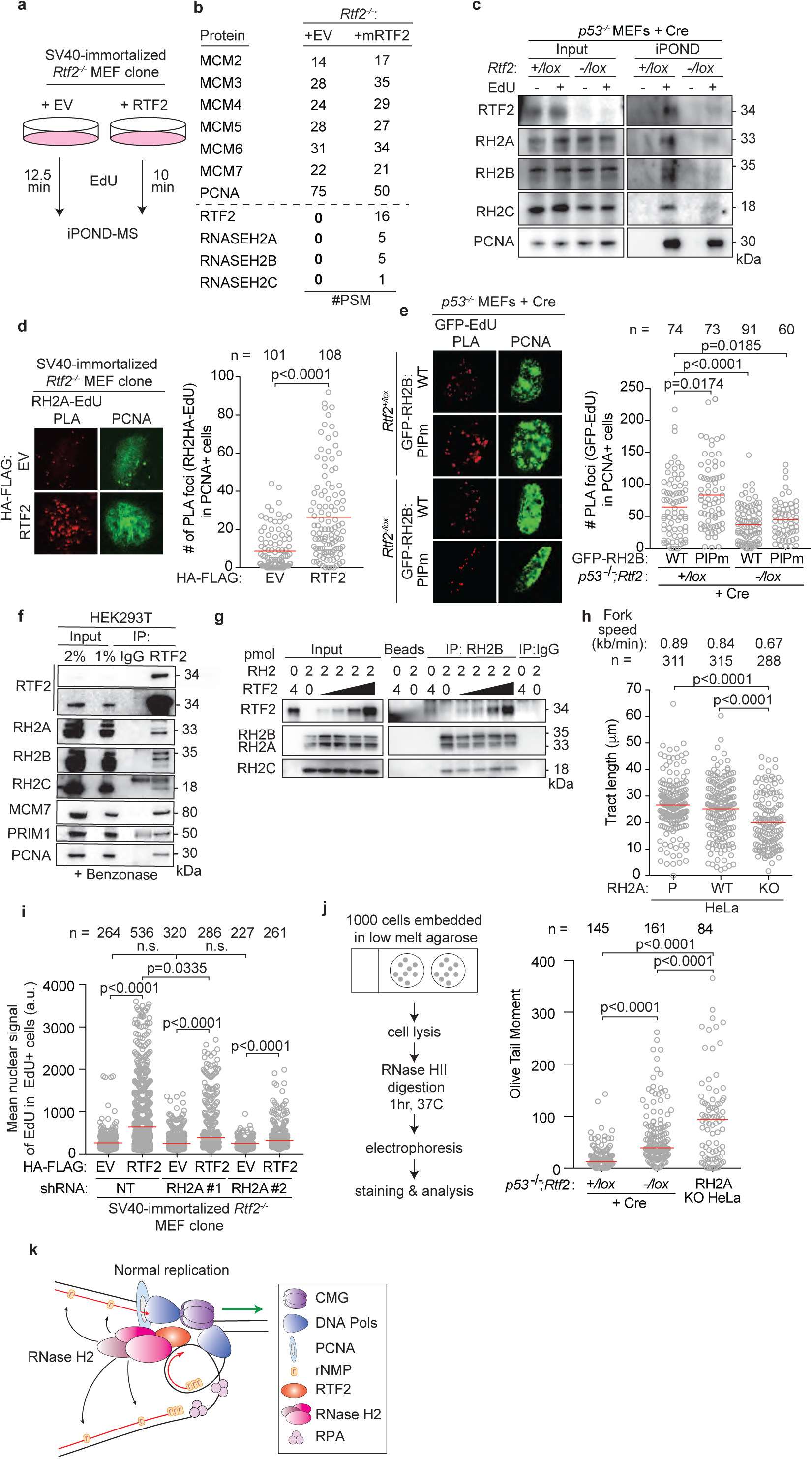
RTF2 recruits RNase H2 to the replication fork to facilitate normal replication and removal of genomic ribonucleotides. **a,** Schematic of the iPOND set-up in SV40-immortlized immortalized *Rtf2^-/-^* MEFs expressing HA-FLAG empty vector (EV) or RTF2 cDNA. EdU pulses were normalized to replication tract lengths. Nascent DNA was purified using streptavidin beads against biotin-conjugated EdU. **b,** Peptide Spectral Match (#PSM) values for indicated proteins from experiment in a. **c,** Representative iPOND immunoblot from *p53^-/-^* MEFs at 72 hrs after Cre. **d,** Left: Representative images of RNASEH2A-EdU nascent proximity ligation assay (nPLA) co-stained with PCNA in SV40-immortalized *Rtf2^-/-^* MEFs expressing HA-FLAG EV or RTF2 cDNA. Right: Quantification of RNASEH2A-EdU foci in PCNA-positive cells. **e,** Left: Representative images from GFP-EdU nPLA co-stained with PCNA in *p53^-/-^* MEFs expressing GFP-tagged wildtype (WT) or PIP box mutant (PIPm) RNASEH2B at 72 hrs after Cre. Right: Quantification of GFP (RNASEH2B)-EdU foci in PCNA-positive cells. Note that endogenous RNase H2 is present in these cells. **f,** Representative immunoblots following immunoprecipitation of RTF2 from HEK293T cells. **g,** Representative immunoblot from immunoprecipitation of recombinant RNase H2 complex and RTF2 expressed in *E. coli*. Protein amount (pmol) indicated above each lane; range of RTF2 is 0.5, 1, 2, 4 pmol. Images for input and pulldown of RTF2 are separate exposures of the same blot. **h,** Quantification of representative experiment of replication tract lengths in CRISPR-edited RNASEH2A knockout (KO) or wildtype (WT) HeLa cells. **i,** Quantification of mean nuclear signal of EdU in EdU-positive SV40-immortalized MEFs, combined from 4 biological replicates. **j,** Left: Schematic of neutral comet assay post RNase HII-digestion. Right: Quantification of olive tail moment in *p53^-/-^* MEFs (72 hrs after Cre), combined from 4 biological replicates. RNASEH2A KO HeLa cells serve as positive control. **k,** Model of RTF2’s function to maintain RNase H2 levels at the replisome during normal replication. Experiments were conducted at least three times in biological replicates with consistent results for c,d,f,g,h,i. Experiment shown in e were conducted twice, in two biological replicates. Mean is shown with red line for d,e,h.i,j. Experiments were blinded prior to analysis for g and h. Average fork speeds are listed above each sample for g. Outliers removed with ROUT (1%) for g. Significance was evaluated by Kruskal-Wallis ANOVA with a Dunn’s post-test. P = Parental, RH2A= RNASEH2A, RH2B= RNASEH2B, RH2C= RNASEH2C, RH2 = RNASEH2 complex, Ctrl = Control.

RNase H2 has been previously reported to localize to the replication fork through its PCNA-interacting protein-box (PIP-box) in the RNASEH2B subunit^13,14^. Using cells overexpressing WT or PIP-box mutated RNASEH2B (RNASEH2B^F300A;F301A^), we asked if RNASEH2B localization to nascent DNA was dependent on the intact PIP box and/or RTF2. nPLA revealed equivalent numbers of PIP-mutated and WT RNASEH2B foci when RTF2 was present (*p53^-/-^;Rtf2^+/lox^*MEFs + Cre). However, the number of nPLA foci diminished when RTF2 was absent (*p53^-/-^;Rtf2^-/lox^* MEFs + Cre), and the extent of the decrease was similar for the WT and PIP-mutant RNASEH2B (Fig. 2e, Extended Data 5g). In combination with the above data showing that endogenous RNase H2 no longer localizes to nascent DNA in the absence of RTF2, we conclude that RTF2 is a major regulator of RNase H2 localization to the replisome. Moreover, RNASEH2B^F300A;F301A^ and PCNA co-immunoprecipitated to the same extent as wildtype RNASEH2B, suggesting that the RNASEH2B-PCNA interactions are by and large indirect (Extended Data Fig. 6a)^13,14^.

To test if RNASEH2A and RTF2 were found in the same complexes in cell, we used endogenously- or exogenously-tagged RTF2 to perform immunoprecipitations and LC-MS analysis. Indeed, they were found to interact (Extended Data Fig. 6b-i), and the interactions were confirmed by co-immunoprecipitation of endogenous proteins (Fig. 2f). Immunoprecipitations of recombinant proteins purified from *E.coli* showed direct interaction between RTF2 and RNase H2 (Fig. 2g, Extended Data Fig. 6j,k)^15^. Together, these data show that RTF2 localizes the RNase H2 complex to nascent chromatin through a direct interaction. Like RTF2 itself, RNase H2 levels at replication forks are dynamically regulated by the DDI proteins.

We next evaluated whether RNase H2 deficiency phenocopies RTF2 deficiency, which would be consistent with RTF2 regulating RNase H2. Like RTF2-deficient cells, RNASEH2A knockout (KO) cells displayed slow replication speed (Fig. 2h). Additionally, *RNASEH2A*^-/-^ ;*p53^-/-^* HCT-116 cells, *RNASEH2A*^-/-^ HeLa cells,^16^ and *RNASEH2A*-depleted BJ cells, exhibited other phenotypes consistent with a replication defect, including slow growth and a significant decrease in mean nuclear signal of EdU (Extended Data Fig. 7a-h)^4,16^. These defects could be rescued by expression of WT RNASEH2A, but not catalytic dead RNASEH2A^D34A;D169A^ or ribonucleotide excision deficient (isolation of function (IOF)) RNASEH2A^P40D;Y210A^ (Extended Data Fig. 7d-f)^17,18^. Expression of RNase H1, another mammalian ribonuclease with nucleolytic activity against RNA-DNA hybrids^19^, could not rescue the growth and EdU defects observed in RNASEH2A KO cells (Extended Data Fig. 7i-k). These data indicate that RNase H2, but not RNase H1, promotes DNA replication in mammalian cells.

Depletion of RNASEH2A in RTF2-deficient cells did not further diminish the mean nuclear signal of EdU (Fig 2i). Furthermore, under those conditions, expression of an RTF2 cDNA could not fully rescue the replication defect, suggesting that RNase H2 and RTF2 function in the same pathway during DNA replication (Fig. 2i, Extended Data Fig. 7l,m). Since RNase H2 removes ribonucleotides embedded in the newly replicated DNA and consequently *p53^-/-^*;*Rnaseh2b^-/-^* MEFs harbor high levels of ribonucleotides in their genome^4^, we asked whether cells without RTF2 show a similar increase in genomic ribonucleotides. Using a neutral comet assay incorporating RNase HII digestion (Fig.2j, left panel), we observed an increased load of genome-embedded ribonucleotides in RTF2-deficient cells as compared to RTF2-competent cells (Fig. 2j, Extended Data Fig. 7n). Alongside increased genomic ribonucleotides, we observed poly-ADP-ribosylation and S phase-specific γH2AX phosphorylation in RTF2-deficient cells, two indicators of replication-dependent DNA damage following inappropriate TOP1-mediated ribonucleotide processing (Extended Data Fig. 7o-s)^4,16,20–23^.

Taken together, our experiments show that failure to localize RTF2 and RNase H2 to the replisome leads to slow replication fork speeds and accumulation of genome-embedded ribonucleotides. These data support a model whereby RTF2 localizes RNase H2 to traveling replication forks to facilitate removal of genome-embedded ribonucleotides during replication (Fig. 2k). Genomic ribonucleotide incorporation may result from inappropriate incorporation by the replicative polymerases, inefficient Okazaki fragment maturation, or replication-associated DNA repair and their removal by RTF2-localized RNase H2 is important for the stability of the genome^24–26^.

### Regulation of RTF2, RNase H2, and PRIM1 coordinate the response to replication stress and replication fork restart

Proper RTF2 levels at the replisome are necessary for the correct cellular response to replication stress^3^. Cells with an overabundance of replisome-associated RTF2 under conditions of DDI1/2-depletion exhibit increased sensitivity, an accumulation of ssDNA, chromosomal breakage, and inefficient replication restart in response to treatment with replication stress-inducing agents^3^. Given that we showed above that RTF2 localizes RNase H2 to the replication fork to promote unperturbed DNA replication, we examined whether RTF2 and RNase H2 function together in the response to replication stress.

DDI1/2-depleted cells exhibit increased sensitivity to replication stress-inducing agents, including hydroxyurea (HU), aphidicolin, mitomycin C, and gemcitabine, that can be rescued by RTF2 depletion^3^. Like RTF2 depletion, knockdown of RNASEH2A rescued the sensitivity of DDI1/2-depleted cells to HU (Fig. 3a). Even in control cells, depletion of RTF2 or RNASHE2A reduced sensitivity to increasing doses of HU (Fig. 3a), suggesting that the overall level of RTF2 and RNASEH2A at the replisome may dictate the response to local replication stress. Depletion of RNASEH2A, like depletion of RTF2, reversed other phenotypes seen in cells without DDI1/2, including elevated p-RPA S4/8 signaling during low dose HU treatment (Fig. 3b, Extended Data Fig. 8a) and genome instability as measured by presence of chromosomal abnormalities, which are a direct outcome of an inappropriate replication stress response (Fig. 3c, Extended Data Fig. 8b,c). These data suggest that DDI1/2-dependent removal of RTF2 and RNase H2 from stressed replication forks is essential for the proper replication stress response.

**Figure 3.**
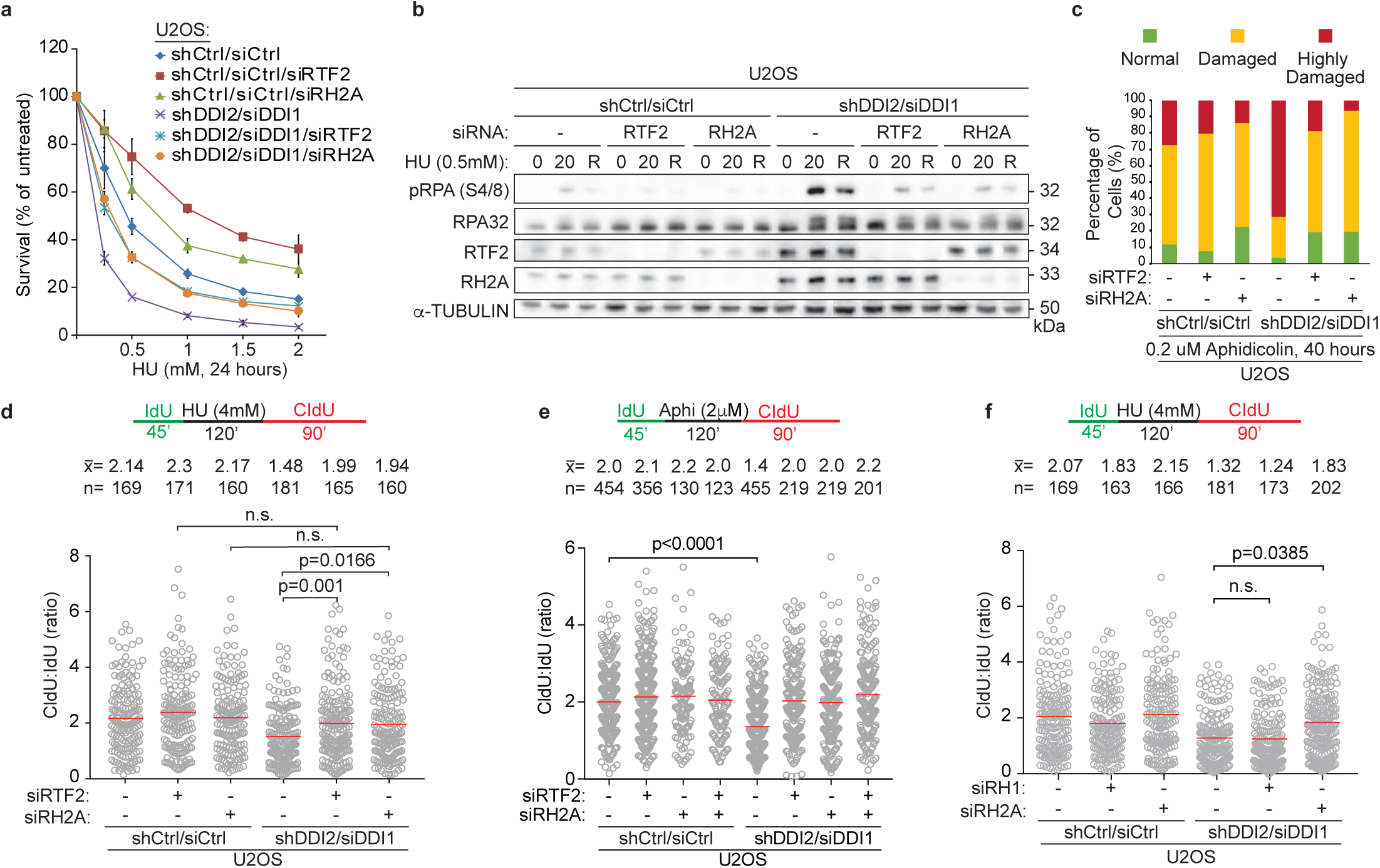
RTF2 and RNase H2 coordinate proper response to replication stress. **a,** Representative cellular survival in HU-treated U2OS cells transduced or transfected with indicated RNAi reagents. **b,** Representative immunoblot in U2OS cells transduced or transfected with indicated RNAi reagents and treated with HU (0 = untreated, 20 = 20 hr treatment, R = 20 hr treatment followed by 8 hr release). **c,** Quantification of chromosome damage in a representative experiment of aphidicolin-treated U2OS cells transduced or transfected with indicated RNAi reagents. **d, e, f,** Quantification of representative experiments of CldU:IdU tract length ratio in DNA combing fork restart assay with fork stalling by HU (d,f) and aphidicolin (e). Experiment was conducted three times in biological replicates with technical triplicates for a. Experiments were conducted at least three times in biological replicates with consistent results for b,c,d,e,f, Mean is shown with red line for d,e,f. Error bars represent standard deviation. Experiments were blinded prior to analysis for c,d,e,f. Average CldU:IdU ratios are listed above each sample for d,e,f. Outliers were removed with ROUT (1%) for d,e,f. Significance evaluated by Kruskal-Wallis ANOVA with a Dunn’s post-test. RH2A= RNASEH2A, RH1 = RNASEH1

A striking phenotype associated with DDI1/2-depletion was a decreased percentage of restarted replication forks and a decreased efficiency of fork restart after release from HU treatment^3^. To further characterize the replication restart defects under these conditions, we used a modified DNA combing assay in which the initial 45 minute pulse of IdU is followed by complete replication block with 4 mM HU (Extended Data Fig. 8d). Following removal of HU, the length of the second nascent DNA tract (CldU) is compared to the length of first tract (IdU) as a proxy for replication restart efficiency (Extended Data Fig. 8d). The fork restart defect observed upon treatment with HU in DDI1/2-depleted cells was rescued to the same extent by depletion of RTF2 or RNASEH2A, suggesting that both RTF2 and RNase H2 must be removed from replication forks to promote replication restart following fork stalling (Fig. 3d). Importantly, RTF2- or RNASEH2A-depleted cells had unperturbed ratios of CldU:IdU when untreated (Extended Data Fig. 8e).

To show that this restart defect was not specific to fork stalling by HU, we attempted to repeat this assay using camptothecin (CPT), gemcitabine, and aphidicolin. Replication could not restart within a short time after complete stalling induced by CPT or gemcitabine (Extended Data Fig. 8f,g). This is most likely due to an inability to directly restart lesions induced by these agents, which necessitate nucleolytic processing and homology-directed repair^27^. However, replisomes completely stalled by 2 μM aphidicolin treatment were able to restart in a timely manner, and fork restart depended on the depletion of RTF2 and RNASEH2A (Fig. 3e). These data show that RTF2 and RNase H2 must be removed from stalled replication forks to promote replication restart. Of note, this effect was specific to RNase H2, as depletion of RNase H1 did not rescue the fork restart defect in DDI1/2-depleted cells (Fig. 3f, Extended Data Fig. 8h,i).

These findings are consistent with the DDI/RTF2 pathway regulating the RNase H2 complex at the replication fork to allow proper recovery from transient DNA replication stress. If persistent, RNase H2 seems to interfere with an activity necessary for efficient replication restart of stalled forks. As RNase H2 processes RNA-DNA hybrids, we hypothesized that persistent RNase H2 endonucleolytically processes an RNA primer necessary to resume replication. To test this hypothesis, we examined the contribution of the two AEP primases, PRIMPOL^28^ and PRIM1, to the efficiency of replication restart after transient replication stalling.

Despite its reported role in fork restart after UV, depletion of PRIMPOL had no effect on the efficiency of replication fork restart after release from HU (Extended Data Fig. 9a-d, in d, compare lanes 1 and 2)^29–31^. Moreover, the rescue of replication restart under conditions of DDI1/2 and RTF2 (or RNASEH2A) co-depletion was not diminished upon PRIMPOL depletion (Extended Data Fig. 9a-d, in d, compare lanes 9, 10, 11, and 12). However, PRIMPOL depletion resulted in an increase in p-RPA S4/8 signal after HU treatment that remained unchanged with co-depletion of DDI1/2 (Extended Data Fig. 9e). These data are consistent with a role for PRIMPOL in filling in gaps behind the replication fork, a pathway that does not seem to involve DDI1/2, RTF2, or RNase H2^32^.

We next examined the role of PRIM1, the catalytic primase subunit of the DNA polymerase α-primase complex, during replication fork restart. Although cells depleted of PRIM1 maintained an unperturbed ratio of CldU:IdU under untreated conditions, they exhibited decreased fork restart efficiency after treatment with HU (Fig 4a, Extended Data Fig. 10a-c), indicating that replication restart is reliant on PRIM1 activity. This dependency was seen regardless of RTF2 or RNASEH2A status (Fig. 4b, Extended Data Fig. 10a-c). Knockdown of PRIM1 induced p-RPA S4/8 in untreated conditions that was exacerbated with HU treatment (Extended Data Fig. 10d). In a parallel experiment, we found that low levels of PRIM1 inhibitor vidarabine-TP (V-TP, 10μM) did not significantly perturb normal replication but resulted in inefficient restart after HU treatment, an effect that could not be overcome by depletion of RTF2 or RNASEH2A (Extended data Fig 10e-g)^33^. These data suggest that PRIM1 catalytic activity is essential for efficient replication restart in mammalian cells.

**Figure 4.**
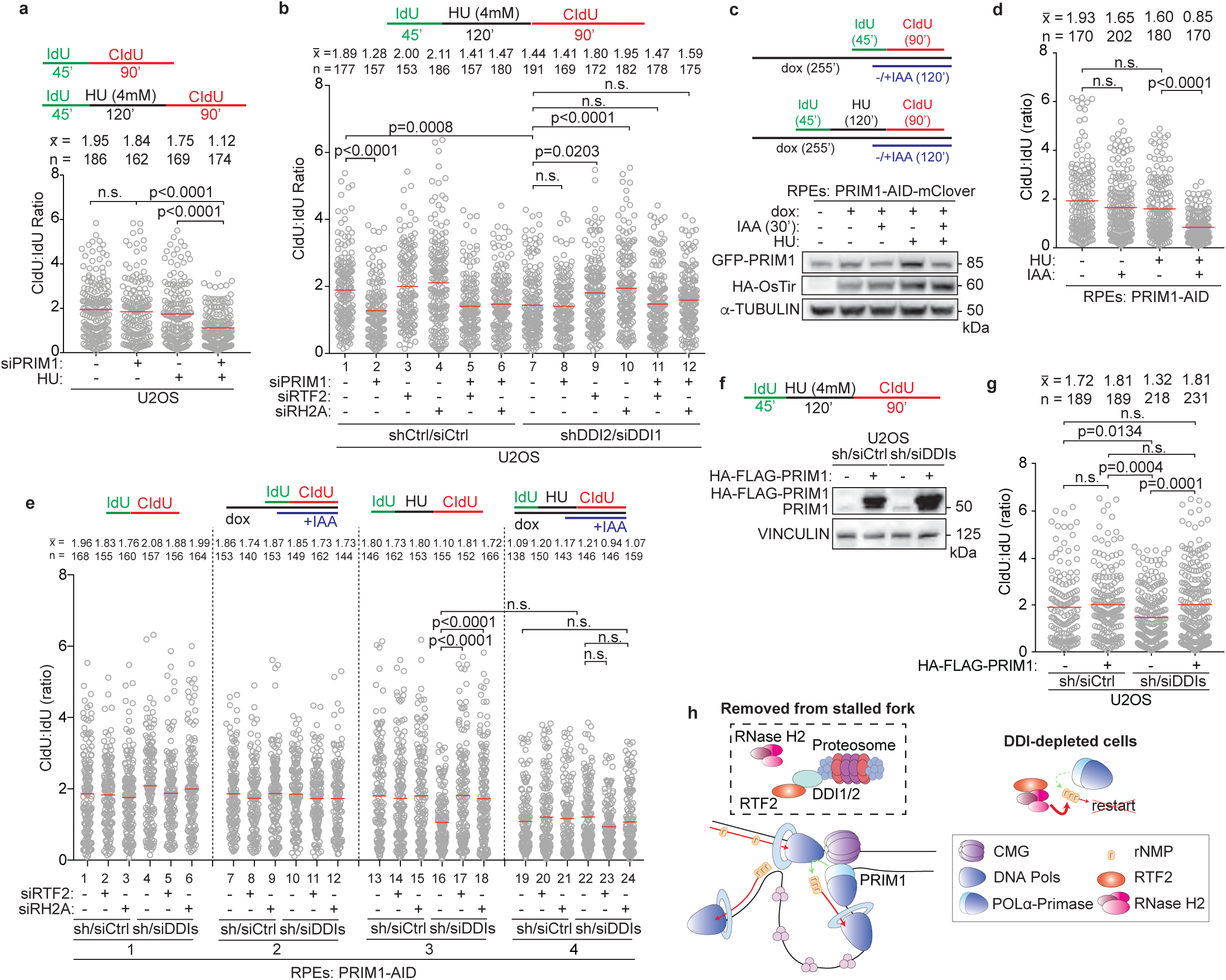
Primase PRIM1 promotes efficient replication restart after stress. **a,b,** Top: Schematic of labeling scheme. Quantification of representative experiment of CldU:IdU tract length ratio in U2OS cells after indicated perturbation. **c,** Top: Schematic of labeling scheme. Dox used to induce expression of HA-OsTir enzyme for targeted PRIM1-AID-mClover degradation. Bottom: Representative immunoblot of PRIM1 levels in endogenously edited PRIM1-AID-mClover RPE cells following indicated treatments. **d,** Quantification of representative experiment of CldU:IdU tract length ratio in cells from experiment in c. **e,** Quantification of representative experiment of CldU:IdU tract length ratio. **f,** Top: Schematic of labeling scheme for experiment in g. Bottom: Immunoblot of HA-FLAG-PRIM1 expression in control and DDI-depleted U2OS cells. **g,** Quantification of representative experiment of CldU:IdU tract length ratio in cells from treatment in f. **h,** Model for removal of RTF2 and RNase H2 from stalled replication forks to allow for restart. Experiments conducted at least three times in biological replicates with consistent results for a,b,c,d,e,g. Mean is shown with red line for a,b,d,e,g. Experiments were blinded prior to analysis for a,b,d,e. Average CldU:IdU ratios are listed above each sample for a,b,d,e,g. Outliers removed with ROUT (1%) for a,b,d,e,g. Significance evaluated by Kruskal-Wallis ANOVA with a Dunn’s post-test. RH2A= RNASEH2A, Ctrl = Control.

As PRIM1’s essential role in DNA replication could confound the results obtained with a 72-hr depletion of PRIM1, an auxin-inducible degron (AID) system was employed to partially degrade PRIM1 within 30 minutes (Fig. 4c)^34^. An acute 20% reduction in PRIM1 expression resulted in a significant decrease in replication restart efficiency (Fig. 4d). The addition of auxin in the untreated condition (no HU) did not significantly change the length of replication tracks in the presence of the first thymidine analog, IdU (Extended Data Fig. 10h). These data indicate that replication restart is exquisitely sensitive to even small changes in PRIM1 levels (Fig. 4d).

To understand the genetic interaction of PRIM1 with the DDI-RTF2 pathway, DDI1/2 were depleted in RPE PRIM1-AID-mClover cells. No further reduction in replication restart efficiency was observed with combined reduction of PRIM1 and DDI1/2 levels (Fig. 4e). However, depletion of PRIM1 prevented the rescue of replication restart seen with the depletion of RTF2 or RNASEH2A in the DDI1/2-depleted setting (Fig. 4e, compare lanes 17,18 to 23,24), suggesting that PRIM1 and RTF2/RNASEH2A function in the same pathway. Depletion of DDI1/2 did not affect the levels of PRIM1 at either progressing or stalled replication forks (Extended Data Fig. 10i). However, DDI1/2-depleted cells overexpressing HA-PRIM1 were proficient in replication restart following stalling by HU, despite retained levels of RTF2-RNase H2 (Fig. 4g, Extended Data Fig. 10j).

Based on the data presented here, we propose that maintaining the balance between RNase H2 and PRIM1 activity at the replication fork allows for repriming and direct replication restart, enabling the resumption of replication without replication fork collapse. While biochemical reconstitution studies in bacteria and yeast have provided evidence for DNA polymerase α-primase-dependent re-priming on the lagging strand,^35–38^ we provide the first evidence of PRIM1-dependent replication fork restart in mammalian cells. Our data show that RTF2, which is dynamically removed from replication forks in the setting of DNA damage, facilitates efficient PRIM1-dependent restart of stalled forks by removing RNase H2 (Fig. 4h).

## Discussion

The identification of DDI1/2 and RTF2 as coordinators of RNase H2 localization to the replication fork implicates the DDI/RTF2/RNase H2 axis as central in regulating both unperturbed DNA replication and the replication stress responses. Our data show that RNase H2 must be tightly regulated during S phase to ensure proper balance between replication repriming and ribonucleotide excision.

We find that localization of RNase H2 to replication forks is RTF2-dependent and PCNA-independent. These findings were surprising at first, given previous reports of PCNA-dependent localization of RNase H2 to sites of replication through RNASEH2B’s PIP box^13^. However they are consistent with prior work identifying PCNA-independent localization of RNase H2 to replication forks through RNASEH2A^13,14^. Further studies will be necessary to identify conditions when the PCNA-RNase H2 interaction is necessary for cellular function.

Future work will also be needed to biochemically and structurally define how DDI1/2 and RTF2 regulate localization of RNase H2 at replication forks to maintain the balance between RNA primer deposition and ribonucleotide excision during unperturbed replication. Under those circumstances, RNase H2 localized by RTF2 might be expected to interfere with PRIM1 priming activity necessary for initiating replication. We suspect that other proteins and regulatory networks will play a role in preventing close association of these proteins while replication is ongoing.

This discovery of RTF2’s role in regulating replication speed and the response to replication stress suggests it is a regulatory hub at the replisome. Fork stalling agents, like aphidicolin and HU, mimic the replication stalling the replisome faces during duplication of difficult-to-replicate regions like common fragile sites and repetitive sequences. Uncovering the mechanisms of RTF2’s regulation will lend insight into the coordination of replication through vast areas of mammalian genome undergoing transient stress.

Both RNase H2 and PRIM1 have been implicated in human disease. *RNASEH2* pathogenic variants have been identified in the inflammatory disease AGS and cancer^39^. A pathogenic variant in *PRIM1* was reported to cause a primordial dwarfism syndrome^40^. Understanding the basic mechanism underlying the balance in their activities during DNA replication will facilitate an understanding of diverse human diseases. Further, the DDI/RTF2/RNase H2/PRIM1 axis contains druggable targets to sensitize cells to replication stress-inducing agents currently entering the clinic.

## Acknowledgements

We thank the members of the Smogorzewska lab for their comments and suggestions during manuscript preparation. A.S. and B.A.C. are grateful to Titia de Lange, Fred Cross, and David Cortez for discussion of this project. We thank the staff of the Proteomics Resource Center (Henrik Molina, Joseph Fernandez, and Brian Dill) for the mass spectrometry analysis presented in the paper. We acknowledge the Transgenic and Reproductive Technology Center (Rada Norinsky), the Comparative Bioscience Center, the Genomics Center (Connie Zhao), the Imaging Center (Alison North) and the Flow Cytometry Center (Svetlana Mazel) and the Cryo-EM Center (Mark Ebrahim) at the Rockefeller University. We thank Titia de Lange, Dan Durocher, and Andrew Jackson for reagents^15^ and cell lines. B.A.C. was supported by the Women and Science Fellowship from The Rockefeller University and by the NRSA Training Grant #GM066699. P.D.R. was supported by the Anderson Center for Cancer Research Fellowship from The Rockefeller University and by F32 GM143866. C.B. and N.J.B. were supported by a Medical Scientist Training Program grant from the National Institute of General Medical Sciences of the National Institutes of Health under award number: T32GM007739 to the Weill Cornell/Rockefeller/Sloan Kettering Tri-Institutional MD-PhD Program. N.J.B was also supported by F30 CA268717. This work was supported by The Gabrielle H. Reem and Herbert J. Kayden Early-Career Innovation Award, R01GM140400, and by Howard Hughes Medical Institute Faculty Scholar Award to AS.

## Contributions

B.A.C. and A.S. designed the study with P.D.R, C.B., N.J.B, and A.S. designing the revision experiments. The original submission was written by B.A.C. and A.S. with input from all co-authors. The revision was written by P.D.R, C.B, B.A.C., and A.S; B.A.C performed many of the experiments, alongside P.D.R. (recombinant and whole cell co-immunoprecipitations, iPOND immunoblots, nascent PLA experiments), C.B. (RTF2 and RH2A co-depletion, comet, combing, and RH1 complementation experiments), and N.J.B. (RNASEH2A complementation experiments and recombinant protein purification). All above authors and A.S. analyzed results. S.S. generated CRISPR-edited 293Ts replacing the *Rtf2* locus with a GFP-AID-hRTF2 cDNA, N.S. generated the CRISPR-edited RPE PRIM1-AID-mClover cells, T.C. analyzed the RNA-sequencing data, F.P.L. performed metaphase spread analysis, M.K. shared unpublished data, and T.W. maintained the *Rtf2* mouse colony and assisted with the mouse experiments, A.S. supervised the study and obtained the funding.

## Methods

### Generation of Rtf2 Mouse Strains, Maintenance and Genotyping

mESCs containing the *Rtf2* gene targeting construct (*Rtf2^tm1a^*(KOMP)*^Wtsi^*) were obtained from the UCDavis KOMP Repository. The Rockefeller transgenics facility injected targeted mESCs into C57BL/6J blastocysts to generate chimeras. *Rtf2^+/stop^* mice were mated with mice expressing Flp^41^ to generate *Rtf2^+/lox^* mice. *Rtf2^+/lox^* mice were then mated with mice expressing Cre recombinase from a ubiquitous EIIa promoter ^42^. All animals were handled according to the Rockefeller University Institutional Animal Care and Use Committee protocols. See supplemental information for summary of mouse strains.

Mouse tail tips were obtained from pups at day 21. Tails were lysed in DirectPCR Lysis Reagent (Mouse Tail) supplemented with 0.2 mg/mL proteinase K according to manufacturer’s protocol. 0.2-1.0 μL of lysate was used for 20 μL PCR reaction. GoTaq**^®^** DNA polymerase master mix was used for PCRs with appropriate primers. Sanger sequencing was performed at GeneWiz. See supplemental information for mESC long range genotyping primers and for mouse genotyping primers.

### Cell Culture

Mouse embryonic fibroblasts (MEFs) were isolated on embryonic day 13.5 from crosses of *Rtf2^+/lox^* and *Rtf2^+lox^* mice (*Rtf2^+/+^, Rtf2^+/lox^*, and *Rtf2^lox/lox^* MEFs) and crosses of *Rtf2^+/lox^* and *Rtf2^+/-^* mice (*Rtf2^+/+^, Rtf2^+/lox^, Rtf2^+/-^*, and *Rtf2^-/lox^* MEFs). MEFs were expanded to obtain cells for early passage primary cell stocks and to immortalize cells with a retrovirus expressing SV40-LT.^43^ Experiments were conducted with littermate MEFs. Multiple MEF lines from individual mothers were isolated and used in replicate experiments. *Rtf2^-/-^* clonal cell lines were also generated from SV40-LT immortalized *Rtf2^-/lox^* MEFs after infection with pMMP Hit & Run Cre recombinase.^44^ For conditional *Rtf2* cell lines, experiments were performed 72-120 hrs after transduction with pMMP Hit & Run Cre recombinase as indicated.^44^ *p53^-/-^; Rtf2^+/lox^* and *Rtf2^-/lox^* MEF lines were generated using CRISPR gene editing at the p53 locus using the pX459 plasmid.^45^ See supplementary information for list of mutagenesis primers and plasmids.

Primary MEFs and BJ foreskin fibroblasts (transformed by HPV16 E6E7 expression and/or immortalized by expression of catalytic subunit of human telomerase (hTERT)) were maintained in DMEM, supplemented with 15% (v/v) fetal bovine serum (FBS), 1X MEM non-essential amino acids, 2 mM L-alanyl-L-glutamine dipeptide, and 100U/mL penicillin-streptomycin. MEFs virally transformed with SV40-LT, HEK 293T cells, U2OS cells (as described in Kottemann *et al*.) and HeLa cells (Jackson lab) were maintained in DMEM supplemented with 10% FBS and the additional supplements described above. RPE *p53^-/-^, pRb^-/-^,* PRIM1-AID-mClover cells (de Lange lab) were maintained in DMEM/F12 supplemented with 10% FBS and the additional supplements described above. HCT-116 *p53^-/-^* cells were maintained in McCoy’s 5A media supplemented with 10% FBS and the additional supplements described above. Upon confluence, cells were dissociated with trypsin and a fraction of the cells were passaged into a new dish. Cells were cryopreserved in their respective media supplemented with 10% DMSO. See supplemental information for summary of cell types.

### Growth and Sensitivity Assays

For growth assays, 2 × 10^4^ - 3 × 10^4^ cells were plated in each well of a 6-well plate in triplicate. Wells were counted on subsequent days using a Z2 Coulter Counter Analyzer (Beckman Coulter). Population doublings were calculated using the following formula: 3.32 x [log(the number of cell harvested)–log(the initial number of cells plated)].

For sensitivity assays, 2.5 × 10^4^ – 4.5 × 10^4^ cells were plated in each well of a 6-well plate in triplicate. The following day drugs were added at indicated concentrations. After 5-6 days in culture, cells were passaged once at appropriate ratios. Cells were counted when untreated wells reached near confluence around 7-9 days. The cell numbers at each dose of drug were divided by the cell number in the untreated sample to calculate the percent survival.

### RNA Preparation, Reverse Transcription, and Real-Time quantitative PCR

Total messenger RNA was extracted from cells using RNeasy Plus Mini Kit. RNA was reverse transcribed using the SuperScript^TM^ III First-Strand Synthesis System. The relative transcript levels of genes of interest were determined by RT-qPCR using Platinum^TM^ SYBR^TM^ Green SuperMix-UDG. All kits were used according to manufacturer’s protocol. Reactions were run and analyzed on Applied Biosystems™ QuantStudio™ 12K Flex system. See supplemental information for RT-qPCR primers.

### Plasmid generation and mutagenesis

cDNA from MEFs or BJ cells was PCR amplified with primers containing attB sites. attB-PCR products were cloned using the Gateway**^®^**system into pDONR223^46^ with BP Clonase II Enzyme Mix. Site directed mutagenesis of pDONR223 derivatives was performed with Agilent QuikChange or Agilent QuikChange kits according to manufacturer’s protocol. pDONR223 derivatives with inserted cDNAs (pENTR vectors) were cloned into destination vectors with LR clonase II Enzyme Mix.^18,47^ Reactions were transformed into chemically competent DH5-α or Stlb3 *E. coli* cells and plated onto Luria-Bertani (LB)/agar plates with the appropriate bacterial selection (kanamycin (50 μg/ml), spectinomycin (50 μg/mL), chloramphenicol (25 μg/ml), or ampicillin (100 μg/ml)). Clones were mini-prepped and sequences were confirmed with Sanger sequencing (GeneWiz). Appropriate clones were maxi-prepped and ethanol precipitated for sterile tissue culture use. See supplemental information for Gateway primers, mutagenesis primers and a list of plasmids.

pEGFP-RNASEH2B was a gift from Andrew Jackson & Martin Reijns (Addgene plasmid # 108697; http://n2t.net/addgene:108697; RRID:Addgene_108697). pMSCVpuro-DEST was a gift from Andrew Jackson & Martin Reijns (Addgene plasmid # 119745; http://n2t.net/addgene:119745; RRID:Addgene_119745). ppyCAG_RNaseH1_WT was a gift from Xiang-Dong Fu (Addgene plasmid# 111906; http://n2t.net/addgene:111906; RRID:Addgene_111906). ppyCAG_RNaseH1_D210N was a gift from Xiang-Dong Fu (Addgene plasmid # 111904; http://n2t.net/addgene:111904 ; RRID:Addgene_111904). pcDNA5-FRT-TO-EGFP-AID was a gift from Andrew Holland (Addgene plasmid # 80075; http://n2t.net/addgene:80075; RRID:Addgene_80075). MLM3636 was a gift from Keith Joung (Addgene plasmid # 43860; http://n2t.net/addgene:43860; RRID:Addgene_43860). pX330-U6-Chimeric_BB-CBh-hSpCas9 was a gift from Feng Zhang (Addgene plasmid # 42230; http://n2t.net/addgene:42230; RRID:Addgene_42230). pGEX6P1-hsRNASEH2BCA was a gift from Andrew Jackson & Martin Reijns (Addgene plasmid # 108692; http://n2t.net/addgene:108692; RRID:Addgene_108692). pMD2.G was a gift from Didier Trono (Addgene plasmid # 12259; http://n2t.net/addgene:12259; RRID:Addgene_12259). psPAX2 was a gift from Didier Trono (Addgene plasmid # 12260 ; http://n2t.net/addgene:12260 ; RRID:Addgene_12260). pLVpuro-CMV-N-EGFP was a gift from Robin Ketteler (Addgene plasmid # 122848; http://n2t.net/addgene:122848 ; RRID:Addgene_122848). pSpCas9(BB)-2A-Puro (PX459) V2.0 was a gift from Feng Zhang (Addgene plasmid # 62988; http://n2t.net/addgene:62988; RRID:Addgene_62988). pMSCV_PM_shRNA_Control_puro was a gift from Steve Elledge^48^. pMMP Hit & Run Cre was a gift from David Livingston^44^. pWZL Cre-hygro was a gift from Titia de Lange^34^. PSKA002 HIS14-SUMO-MCS Expression Vector and PSKA008 HIS14-GFP-MCS-Expression Vector were gifted from Sebastian Klinge.

### Transductions

cDNAs were delivered by retroviral or lentiviral transduction after packaging in HEK 293T cells. 5x10^6^ were plated the evening before transfection. DNA and viral packaging vectors were transfected into cells with TransIT-293 transfection reagent according to the manufacturer’s protocol. The media was changed the next day and after 24 hrs, viral supernatants were harvested and filtered (0.45 μM). Harvests were repeated every 12 hrs for 2 days. Target cells were infected with viral supernatants supplemented with 4 μg/mL polybrene. Stably expressing cells were selected with the appropriate agent ((puromycin (0.5-2 μg/ml), hygromycin (100 μg/ml), blasticidin (600 μg/ml), neomycin (600 μg/ml)). See supplementary information for the list of plasmids.

### Generation of GFP-AID-RTF2 endogenously tagged 293Ts

The pUC19-EGFP-AID-hRTF2 donor construct was generated with classical and In-Fusion**^®^**HD cloning techniques. hRTF2 was amplified with BamHI and NotI digestion sites and ligated into the pcDNA5-FRT-TO-EGFP-AID backbone. Using this as a PCR template, a GFP-AID-hRTF2 construct was amplified with primers compatible for In-Fusion. 5’ and 3’ UTR regions of RTF2 were also amplified with primers compatible with In-Fusion. Subsequently purified PCR products from these reactions were cloned into a HindIII- and EcoRI-digested pUC19 backbone with In-Fusion**^®^**, generating a donor construct containing homolog arms in the 5’ UTR and 3’ UTR and a cDNA for GFP-AID-RTF2. HEK293Ts were transfected using TransIT-293 with this donor construct, a pX330 guide RNA-Cas9 construct, and a single guide RNA construct in a 1:1:1 ratio.^49^ Cells were grown for 48-72 hrs before sorting for GFP+ populations. GFP+ populations were sub-cloned for single cell clones. These clones were verified for integration of the GFP-AID-RTF2 cDNA into the endogenous *Rtf2* locus. See supplemental information for the list of CRISPR cloning primers.

### Immunoblotting

If cells were counted upon collection, an equal number of cells were lysed by resuspension in an equal volume (100 μL per 1x10^6^ cells) of hot Laemmli buffer (Bio-Rad). If cells could not be counted upon collection, whole cell lysates were prepared by lysing cells in Laemmli buffer (4% SDS, 20% glycerol, 0.125 M Tris-HCl (pH=6.8)), to determine protein concentration by Lowry protein assay prior to addition of 2-mercaptoethanol and bromophenol blue. Samples were either sonicated or passed through a tuberculin needle 10 times. Subsequently, samples were boiled for 5 minutes. Equal amounts of protein were separated by sodium dodecyl sulfate polyacrylamide gel electrophoresis (SDS-PAGE) on precast 4–12% Bis-Tris gels. Membranes were blocked for 1 hr in 5% milk in TBST (10 mM Tris-HCl (pH=7.5), 150 mM NaCl, 0.1% Tween 20) and incubated in primary antibodies for 2 hrs at room temperature or overnight at 4°C. Membranes were sufficiently washed in TBST (3x 10 minutes) before being incubated with horseradish peroxidase (HRP)-conjugated secondary antibodies for 1 hr at room temperature, sufficiently washed again and detected by enhanced chemiluminescence. Membranes were visualized with either ImageQuantLAS 4000 or Azure c300 imaging systems. See supplemental information for the list of antibodies.

### siRNA Transfections

Cells were transfected with pools of 3 siRNAs against DDI1 as previously published.^3^ For PRIM1 and PRIMPOL, pools of 3 siRNAs were used. For RNASEH2 and RTF2 depletion, a single siRNA was used. Cells were transfected using Lipofectamine™ RNAiMAX Transfection Reagent according to manufacturer’s instructions with the exception that siRNA-lipid complexes were added to the well and then cells were seeded on top of complexes. Knockdown was measured by RT-qPCR or western blot. See supplemental information for RNAi sequences.

### Immunofluorescence and Nascent Proximity Ligation Assay

#### EdU staining

Cells were pulsed with 10μM EdU for 1 hr, then washed in 1x PBS and fixed in 3.7% (v/v) formaldehyde in PBS at RT for 10 minutes. Cells were washed and permeabilized in 0.5% (v/v) Triton X-100 in PBS. Cells were blocked in either 5% (v/v) FBS in PBS or 3% BSA (w/v) in PBS at RT for 30 minutes. Cells were stained with Click-iT™ EdU Alexa Fluor™ 488 Imaging Kit (Invitrogen, C10337) according to manufacturer’s protocol. Cells were washed and embedded on glass slides with DAPI Fluoromount-G (SouthernBiotech). Mean nuclear signal is mean gray value calculated with image analysis using FIJI.

#### γH2AX staining

Cells were fixed and permeabilized as above. Slides were incubated with primary antibodies in blocking buffer for 2 hrs at room temperature or overnight at 4°C. Cells were washed with blocking buffer and then incubated with secondary antibody. Cells were washed and embedded on glass slides as above. Mean nuclear signal is mean gray value calculated with image analysis using FIJI.

#### Nascent PLA staining

This assay was optimized to detect the amount of protein at the replication fork by labeling with short pulses of EdU. The duration of EdU pulse was determined by DNA combing experiments to label equal amounts of nascent DNA across different conditions. Cells were washed with PBS, permeabilized with 0.5% Triton X-100 in PBS, fixed with 3% formaldehyde/2% sucrose in PBS and blocked with 3% BSA in PBS. EdU was then biotin-clicked and coverslips were incubated with either mouse anti-biotin and rabbit anti-RNASEH2A, or with rabbit anti-biotin and mouse anti-GFP overnight. The next day, antibody-coupled sense and anti-sense probes were used to detect the light chains of rabbit and mouse IgG, respectively, followed by the PLA reaction (DuoLink) according to the manufacturer’s protocol. If the probes are within 30-40 nM of each other, they are ligated and amplified to produce a fluorescent signal. Where indicated, cells were co-stained with PCNA prior mounting on slides. Slides were imaged (Inverted Olympus IX-71 DeltaVision Image Restoration Microscope (Applied Precision) or Axio Observer.A1 fluorescence microscope (Carl Zeiss), equipped with a Plan-Apochromat 63× NA-1.4 oil objective, the AxioCam CCD camera, and the AxioVision Rel Version 4.7 software. Foci were counted using Cell Profiler software using either DAPI or PCNA co-stain to detect nuclei.

### Cell Cycle

Exponentially growing cells were labeled with EdU 1 hr prior to collection and fixation. Cell cycle preparation was performed with Click-iT™ EdU Alexa Fluor™ 647 Flow Cytometry Assay Kit per the manufacturer’s protocol. Total DNA content was stained with FxCycle™ Violet. Stained cells were analyzed on a BD Accuri^TM^ C6 or BD^TM^ LSR II cytometers. Data was analyzed with FlowJo software.

### RNA sequencing and analysis

Total messenger RNA was extracted using RNeasy Plus Mini Kit and DNase treated prior to submission to Rockefeller University’s genomic core for library preparation. Libraries for intron retention were prepared using rRNA depletion and run on NextSeq 500 High Output for 75 base pair paired end reads. Libraries for transcript analysis were prepared using standard Illumina sequencing primers and run on NextSeq 500 High Output for 75 base pair single reads. For intron retention analysis, raw reads were aligned to the mouse genome (mm10) with Stand NGS’s software. Percentage of reads mapping to introns was from determined post-alignment statistics. For mRNA transcript analysis, raw reads were aligned to the mouse genome and analyzed by Tom Carroll (Rockefeller University Bioinformatics) using the NGSpipeR package. Differential expression was determined with DESeq2.

### iPOND

Cells were pulsed with 10μM EdU for differential times to yield similar labeled lengths. The duration of EdU pulse was determined by DNA combing experiments to label equal amounts of nascent DNA across different conditions. iPOND was performed as previously described^11^ with Dynabeads™ MyOne™ Streptavidin C1 (Invitrogen, 65001). Eluate was run only 1 cm into a 4%– 12% Bis-Tris gel (Invitrogen) and submitted for mass spectrometry analysis at Rockefeller University’s Proteomics Core. For iPOND immunoblots, the eluate was run on 4%–12% Bis-Tris gels (Invitrogen) and immunoblotted as indicated.

### Co-immunoprecipitations

Endogenous co-immunoprecipitations were seeded the previous day at a concentration of 10-20 x 10^6^ cells per 15 cm. Cells were treated 1 hr prior to collection with 10 μM MG-132. Cells were then washed 1x with cold PBS, harvested by scraping, and spun at 500 x g for 5 min at 4°C. Cells were subsequently washed 2x with cold PBS. Cells were lysed in 1mL of cold lysis buffer (50 mM HEPES pH=7.5, 150 mM NaCl, 2 mM MgCl2, 0.1% Tween-20, 1x Phosphatase inhibitor cocktail II, 1x protease inhibitor (Roche, cOmplete EDTA-free, 11697498001), 2 mg/mL N-ethylmaleimide). Cells were sonicated (3x 10A for 15 seconds) and treated with benzonase for 30 minutes. Lysates were centrifuged to remove debris. The cleared lysate was incubated with antibodies and Dynabeads™ (Invitrogen) or antibody-coupled Dynabeads™ M-270 Epoxy resin (Invitrogen). Normal IgG was used for antibody controls. Lysates were incubated at 4°C for 2 hrs on nutator and then washed with lysis buffer. Samples were heated for 10 minutes in 2 x LDS buffer to elute and run on 4-12% Bis-Tris SDS-PAGE gel. Membranes were subject to standard immunoblotting detection methods.

### Live cell imaging

Cells were infected with retrovirus carrying a GFP-H2B cDNA. Cells were infected with Cre, seeded in 35mm MatTek (MatTek, P35G-1.5-14-C) dishes and imaged every 10 minutes using Olympus CellVoyager (Olympus, CV1000).

### DNA Combing

Exponentially growing cells were labeled with IdU (100 μM) followed by and CldU label (100 μM). Cells were collected and washed in PBS. Cells were resuspended in 45 μL Resuspension Buffer (PBS) with 0.2% NaN3. An equal volume of 2% low melting agarose Mb grade (BioRad) melted in resuspension buffer was equilibrated at 55°C and added to cells to make agarose plugs. Agarose plugs were injected into digestion buffer (1mg/mL proteinase K, 1% N-Lauroylsarcosine, 0.2% sodium deoxycholate, 100 mM EDTA, 10 mM Tris-HCl, pH 7.5) and incubated overnight at 55°C. Plugs were washed for at least 3 x 1 hr each in TE 1X pH 8.5 with 100 mM NaCl before melting in 1mL combing buffer at 68°C for 20 minutes. After melting, tubes were transfer to 42°C heat block. After 10 minutes, 1 μL beta-agarase (New England Bio Labs) was added without mixing and incubated overnight at 42°C. DNA was poured into Disposable DNA Reservoirs (Genomic Vision) and combed onto silanized coverslips (CombiCoverslips, Genomic Vision or made in house) using the Molecular Combing System (Genomic Vision). Slides were dried for 2 hrs at 65°C. Slides were denatured with 0.5 M NaOH + 1 M NaCl for 8 minutes, followed by dehydration in 70%, 90% and 100% ethanol. Slides were blocked for 1 hr in 5% FBS in PBS. Slides were incubated with primary antibodies overnight at 4°C or 2 hrs at RT, washed and incubated with secondary antibody for 1 hr at RT. Slides were mounted with Fluoromount-G (SouthernBiotech, 0100-01). Fibers were imaged (Inverted Olympus IX-71 DeltaVision Image Restoration Microscope (Applied Precision)) and measured using FIJI software.

For PRIM1-AID experiments, cells were pulsed with indicated drugs for the following times: doxycycline for 4 hrs and 15 minutes, IdU for 45 minutes, 4 mM HU for 120 minutes, CldU for 90 minutes, IAA for 120 minutes.

### Silanization of Coverslips

Coverslips were prepared as previously published^50^ with modification of the plasma cleaning step, wherein we used Gatan Model 950 Advanced Plasma System for 10 minutes with atmospheric air.

### Metaphase Spreads

Cells were treated for 40 hrs with aphidicolin. In the final 90 minutes of treatment, cells were co-incubated with colcemid (0.1 μg/mL). Cells were harvested and incubated in 5 mL 0.075 M KCL for 10 minutes before being fixed with the addition of 1 mL methanol and acetic acid (3:1). Cells were resuspended in 10 mL of methanol and acetic acid (3:1) overnight at 4°C. Cells were dropped onto wet slides and dried at 42°C for at least 1 hr before staining with 8% (v/v) KaryoMAX^TM^ Giemsa in 1x Gurr buffer for three minutes, washing in 1x Gurr buffer for 3 minutes, washing in water for 3 minutes, and drying. Dry slides were then imaged on the Metasystems Metafer slide scanning platform.

### RNase HII-digested Neutral Comet Assay

Neutral comet assays were performed according to manufacturer’s protocol (Trevigen) with the addition of an RNase HII digestion step. Cells were harvested 72 hrs after Hit & Run pMMP Cre retroviral infection.^44^ Following cell lysis, slides were flooded with 1 X ThermoPol Buffer (NEB) for 5 minutes at 25°C to equilibrate and digested with 10U RNase HII for 1 hr at 37°C. To measure DNA breaks following RNase HII digestion, slides were stained with SYBR Green (1:10,000) in PBS. Comet tail moments were visualized or Axio Observer.A1 fluorescence microscope (Carl Zeiss), equipped with a Plan-Apochromat 40× NA-1.4 oil objective, the AxioCam CCD camera, and the AxioVision Rel Version 4.7 software and scored with OpenComet software.^51^

### Recombinant Protein Purification and Co-immunoprecipitation Experiments

hRTF2 and the hRNASEH2BCA (RNASEH2B is GST tagged) complex were purified from *E.coli* using standard IMAC and GST pulldown, respectively, followed by further purification using an AKTA™ FPLC. hRTF2 and RNaseH2 complex were incubated together at 4°C for 2 hrs in mixing buffer (20 mM HEPES, pH=7.5, 1 mM DTT, 0.01% NP-40, 5% glycerol, 0.1 mg/mL BSA, 1x cOmplete protease inhibitor, 50 mM NaCl; 50uL total reaction). Benzonase was added at final concentration of 1 uL per 1 mL of mixing buffer. After mixing, an additional 1uL of 5M NaCl was added to each reaction to increase final NaCl concentration to 150 mM. Dynabeads™ (Invitrogen) were added to each reaction and incubated 4°C for 1 hr. Beads were washed with 1 mL wash buffer (20 mM HEPES, pH=7.5, 1 mM DTT, 0.01% NP-40, 5% glycerol, 1x cOmplete protease 126 inhibitor, 150 mM NaCl) for 5 minutes. Washes were repeated four times before proteins were eluted in 20uL Laemmli buffer at 100°C for 5 minutes. Lysates were subjected to standard immunoblotting techniques. Sheep polyclonal anti-human RNase H2 complex antibody was a gift from Andrew Jackson & Martin Reijns.^15^

### Quantification and Statistical Analysis

All ANOVA and t-test analysis was done using Graphpad Prism software. Image quantification was completed with FIJI software. For DNA combing restart assays, outliers were identified and removed with ROUT (1%).

---

**Extended Data Figure 1.**
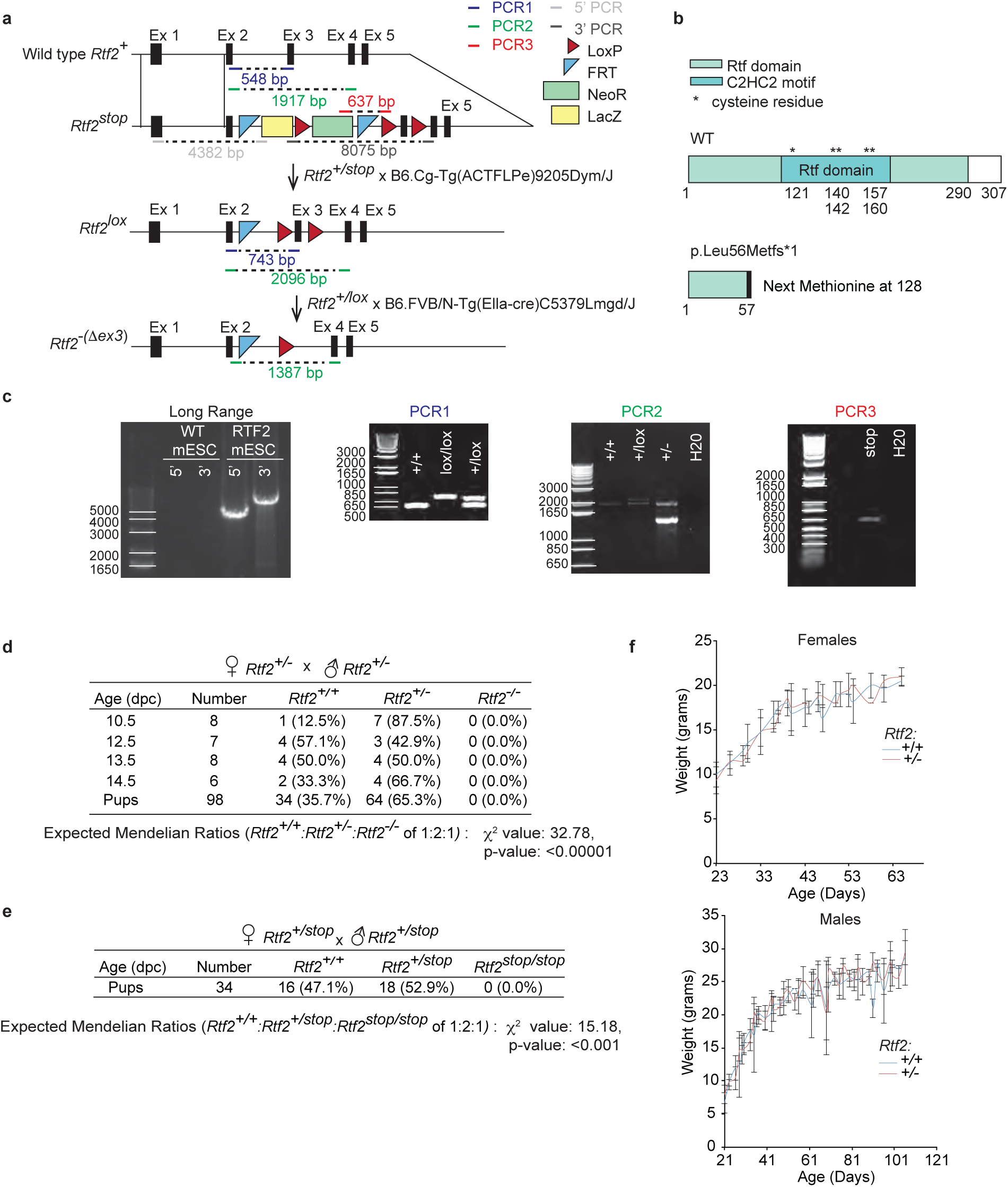
RTF2 is necessary for *in vivo* viability. **a,** Schematic of crosses to generate the genetic knockout of RTF2 in *Mus musculus*. Mouse embryonic stem cells (mESCs) containing an *Rtf2^tm1a(KOMP)Wtsi^ (Rtf2^stop^)* allele (Knockout Mouse Project [KOMP]) were injected into mouse blastocysts (B6(Cg)-Tyr^c-2J^/J, Jackson Labs) to create chimeric mice, which through appropriate crosses generated *Rtf2^+/stop^* mice. *Rtf2^+/stop^* mice were bred with mice expressing Flp recombinase and subsequently mice expressing Cre recombinase under a ubiquitous promoter. These crosses induced the loss of exon 3 to yield *Rtf2^+/Δexon3^* pups. The loss of exon 3 will herein be referred to as *Rtf2^-^*. *Rtf2* mice were maintained on C57BL/6 background. PCR primers and product sizes are indicated on the *Rtf2* alleles. **b**, Schematic of WT RTF2 protein. Loss of exon 3 results in early truncation of RTF2. **c**, Representative genotyping PCR products run on 0.8% agarose gel and stained with EtBr. Long range PCR products amplified from the KOMP mESCs confirmed the presence of the FRT and LoxP sites after excision and analysis with Sanger sequencing. PCR1, PCR2, and PCR3 products amplified from DNA extracted from mouse tail tips were used to confirm genotypes. **d,** Genotypes from litters of *Rtf2^+/-^* female mice crossed with *Rtf2^+/-^*male mice showing embryonic lethality. **e,** Genotypes from litters of *Rtf2^+/stop^*female mice crossed with *Rtf2^+/stop^*male mice. *Rtf2^stop/stop^* mice showing embryonic lethality. **f,** Weights from *Rtf2^+/-^* and *Rtf2^+/+^* mice. For each point, n≥2 mice. n for given genotypes, female *Rtf2^+/+^*= 15, female *Rtf2^+/-^* = 21, male *Rtf2^+/+^* = 14 and, male *Rtf2^+/-^* = 34. Chi-squared test statistic (X^2^) and p-values are indicated for pups. Error bars indicate standard deviation.

**Extended Data Figure 2.**
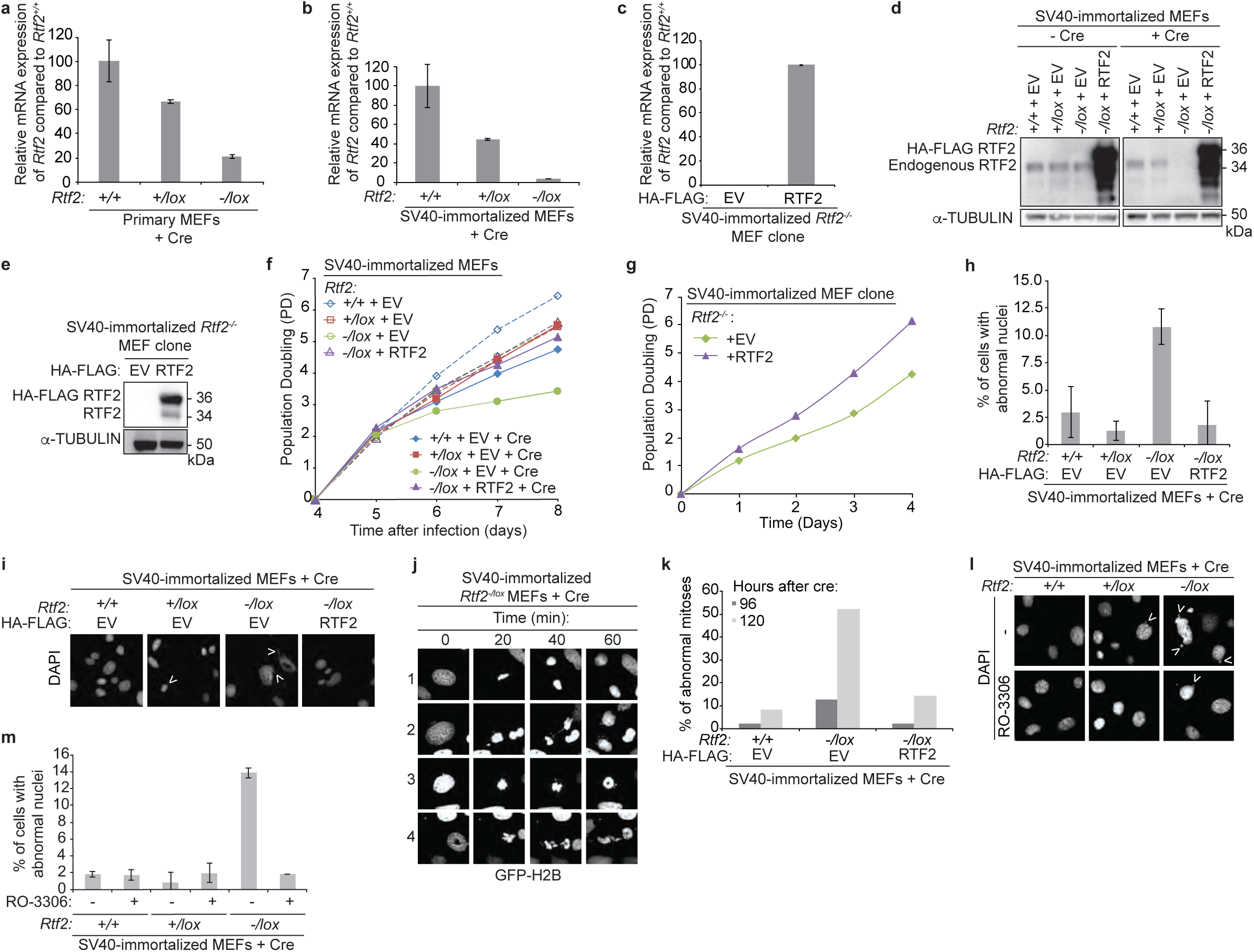
Phenotypes of RTF2-deficient SV40-immortalized MEFs and clones are consistent with phenotypes observed in primary MEFs. **a,b,c**, Representative RT-qPCR of relative *Rtf2* mRNA transcript levels of cDNA prepared from primary MEFs (a) or SV40-immortalized (b) MEFs transduced with Hit & Run Cre recombinase retrovirus 96 hrs before harvest, or from an SV40-immortalized MEF *Rtf2^-/-^* clone expressing empty vector (EV) or RTF2 cDNA (c)^44^. Expression was normalized to β*-actin* expression. **d**, Representative immunoblot of whole cell lysates for RTF2 deletion in *Rtf2^-/lox^* SV40-immortalized MEFs expressing empty vector (EV) or HA-FLAG-mRTF2 (RTF2) cDNA constructs transduced with pWZL Cre-hygro retrovirus 120 hrs before harvest. α-tubulin represents loading control. **e**, Representative immunoblot of whole cell lysates in SV40-immortalized RTF2-deficient sub-cloned MEF lines expressing empty vector (EV) or HA-FLAG-mRTF2 (RTF2) cDNA constructs. α-tubulin represents loading control. **f, g**, Representative growth curves of MEFs 72 hrs after transduction with Hit & Run pMMP Cre (f) or of SV40-immortalized RTF2-deficient sub-cloned MEF lines expressing empty vector (EV) or HA-FLAG-mRTF2 (RTF2) cDNA constructs (**g**)^44^. **h**, Quantification of percentage of cells with abnormal nuclei based on DAPI staining from (i). i, Representative images of DAPI staining from indicated SV40-immortalized MEFs expressing empty vector (EV) or HA-FLAG-mRTF2 (RTF2) cDNA constructs and transduced with pWZL Cre-hygro retrovirus 120 hrs before imaging. Arrows indicate abnormal nuclei. j, Representative images of GFP-H2B staining in live *Rtf2^-/-^* SV40-immortalized MEFs expressing empty vector (EV) and transduced with pWZL Cre-hygro retrovirus 120 hrs prior to analysis undergoing mitosis. In row 1, a cell enters mitosis and undergoes a normal, efficient division. In rows 2-4, cells fail to complete successful mitosis, resulting in lagging chromosomes. k, Quantification of abnormal mitoses from live-cell imaging of *Rtf2^-/-^* SV40-immortalized MEFs expressing GFP-H2B and either empty vector (EV) or HA-FLAG-mRTF2 (RTF2) cDNA constructs and transduced with pWZL Cre-hygro retrovirus as in (**j**). l, Representative images of DAPI-stained nuclei from SV40-immortalized MEFs transduced with pWZL Cre-hygro retrovirus for 120 hrs and then treated with CDK1 inhibitor (RO-3306) for an additional 24 hrs to prevent mitosis. Cells fixed and imaged 144 hrs after Cre retroviral transduction. Arrows indicate abnormal nuclei. m, Quantification of abnormal DAPI-stained nuclei in images shown in (**l**). Experiments were conducted at least three times in biological replicates with consistent results for d,e,f,g,h,i. Averages from two biological replicates plotted in m. Error bars represent standard deviation. Significance was evaluated by Kruskal-Wallis ANOVA with a Dunn’s post-test.

**Extended Data Figure 3.**
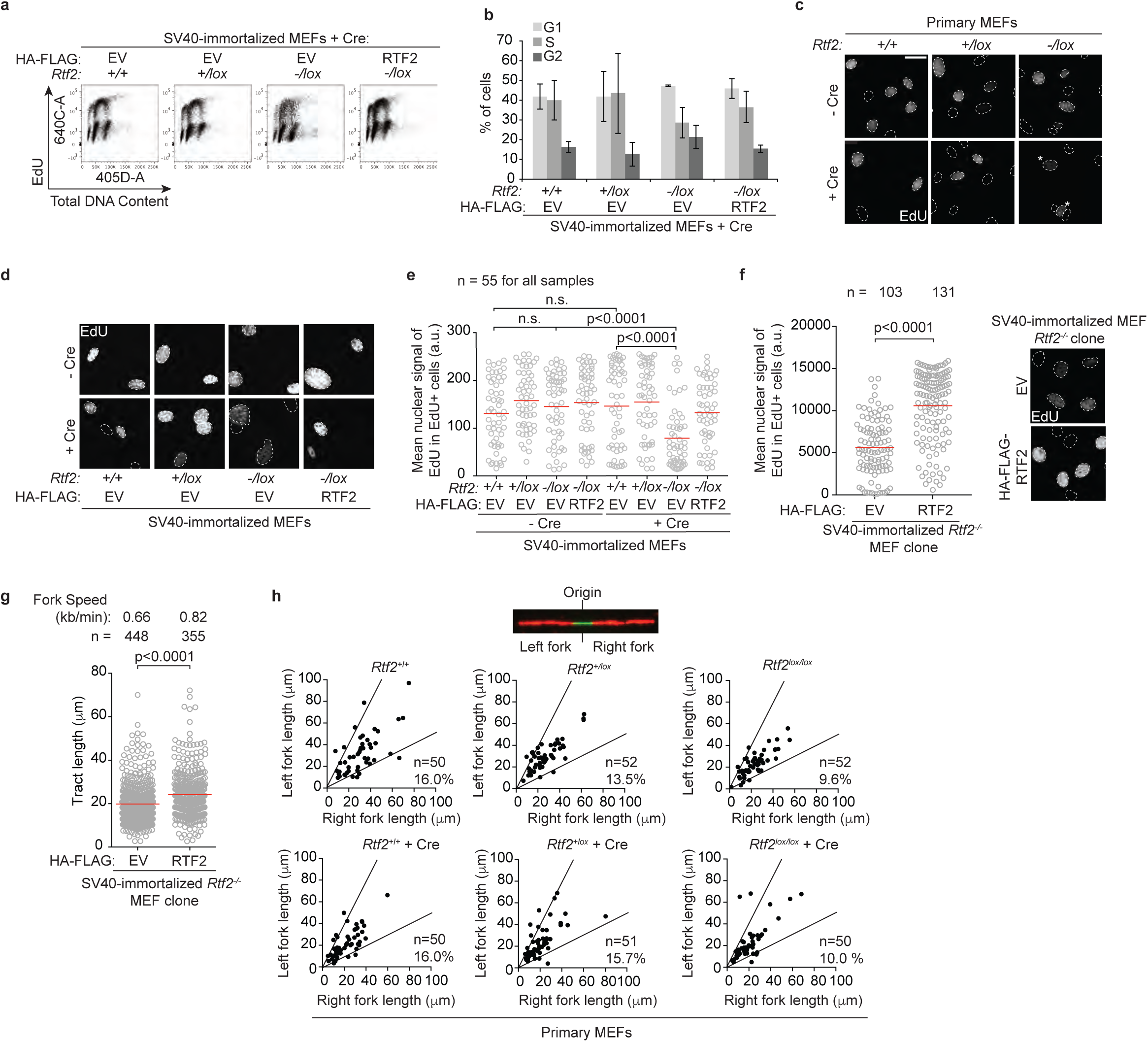
S-phase and replication defects in the absence of RTF2 are not due to replication fork instability. **a,** Representative cell cycle profiles from flow cytometry of indicated SV40-immortalized MEFs transduced with pWZL Cre-hygro retrovirus 120 hrs before analysis. *Rtf2^-/-^* cells incorporate less EdU at a comparable total DNA content in mid-S. **b,** Average percentage of G1, S phase, and G2 gated populations from a. **c,** Representative immunofluorescence of primary MEFs after infection with Hit & Run Cre 72 hrs before fixation.^44^ Cells were pulsed with EdU for 1 hr prior to fixation. Nuclei are outlined in dashed lines based on DAPI staining. Asterisks indicate EdU-positive cells in the *Rtf2^-/lox^* + Cre sample. Quantification is shown in Fig. 1d. **d,** Representative immunofluorescence of SV40-immortalized MEFs expressing empty vector (EV) or HA-FLAG-mRTF2 (RTF2) cDNA constructs and transduced with pWZL Cre-hygro retrovirus 120 hrs before fixation. Cells were pulsed with EdU for 1 hr prior to fixation. Nuclei are outlined in dashed lines based on DAPI staining. **e,** Quantification of mean nuclear signal of EdU from EdU-positive cells from d. Each dot represents one EdU-positive cell. **f,** Left: Quantification of representative experiment of mean nuclear signal of EdU from EdU-positive SV40-immortalized *Rtf2^-/-^* sub-cloned MEFs expressing empty vector (EV) or HA-FLAG-mRTF2 (RTF2) cDNA constructs. Each dot represents one EdU-positive cell. Right: Representative immunofluorescence images of MEFs. Cells were pulsed with EdU for 1 hr prior to fixation. Nuclei are outlined in dashed lines based on DAPI staining. **g,** Quantification of representative experiment of replication tract lengths of progressing fork species in SV40-immortalized *Rtf2^-/-^* sub-cloned MEFs expressing empty vector (EV) or HA-FLAG-mRTF2 (RTF2) cDNA constructs. **h,** Top: Schematic of replication initiation sites, identified as species where the second label (CldU) flanks the first label (IdU). Left fork length is plotted against right fork length to determine fork symmetry. Bottom: Representative experiment showing fork symmetry from primary MEFs 72 hrs after transduction with Hit & Run Cre^44^. Left fork length is plotted against right fork length. Lines represent arbitrary cutoffs for replication forks with symmetry less than 2 and greater than 0.5. Percentages in the bottom right corner represent the percentage of asymmetric forks. Experiments were conducted at least three times in biological replicates with consistent results for a,b,c,d,e,f,g,h. Error bars represent standard deviation. Mean is indicated with a red line for e,f,g. Experiments were blinded prior to analysis for c,d,e,f,g. Average fork speed listed above each sample for g. Outliers removed with ROUT (1%) for g. Significance evaluated by Kruskal-Wallis ANOVA with a Dunn’s post-test.

**Extended Data Figure 4.**
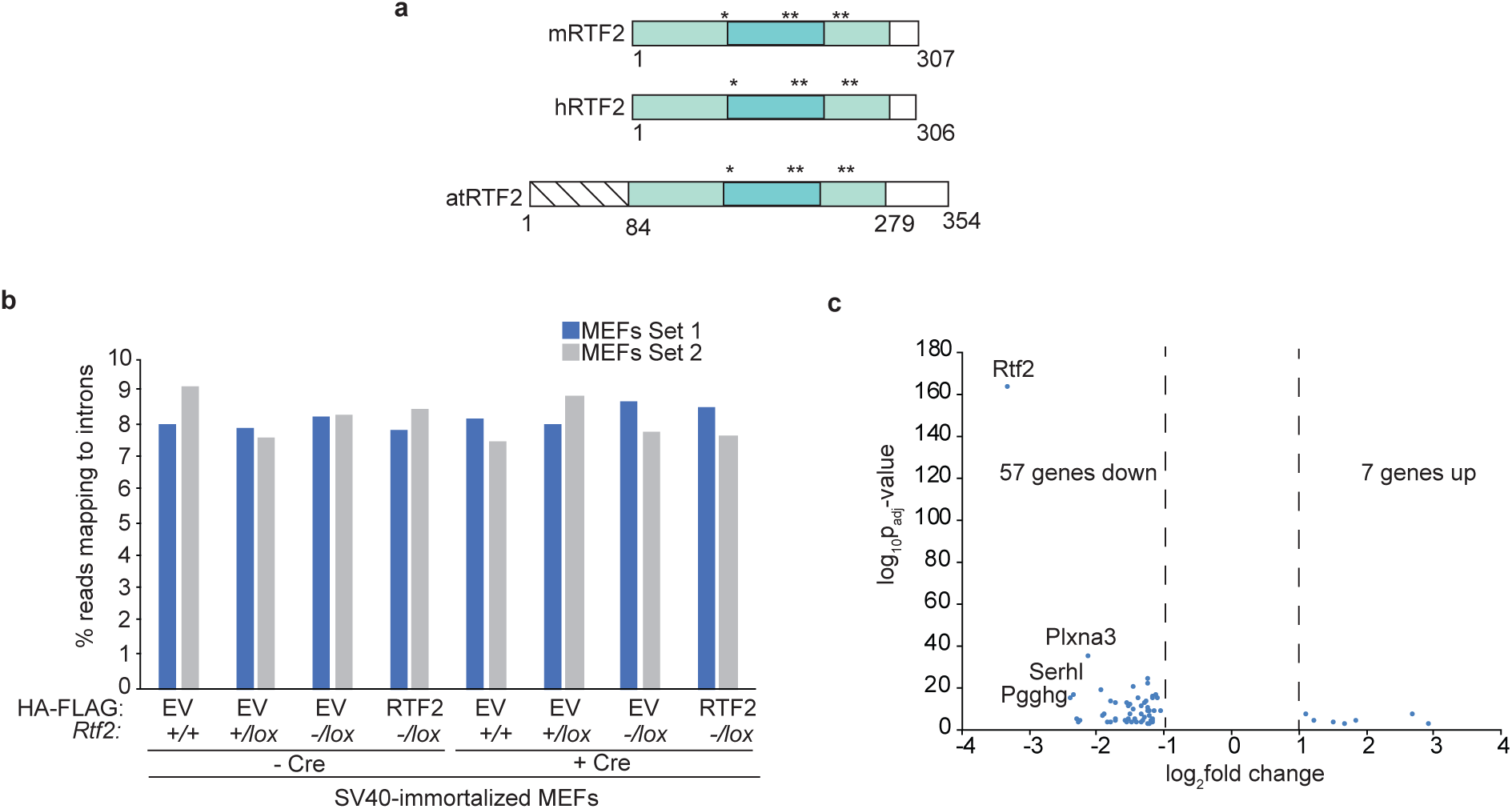
RTF2-deficient MEFs do not display changes in global intron retention or gene expression. **a,** Schematic of RTF2 from *Mus musculus, Homo sapiens* and *Arabidopsis thaliana*. Amino acids 7-63 in atRTF2 are implicated in an intron-retention defect in plants. **b,** Percent of reads from paired-end RNA-seq mapping to introns. Genotypes are indicated for SV40-immortalized MEFs expressing cDNA for HA-FLAG-empty-vector (EV) or HA-FLAG-mouse-RTF2 (RTF2) 120 hrs after transduction with pWZL Cre-hygro retrovirus **c,** Volcano plot showing log_10_p-values against log_2_fold change for the significant differentially expressed genes as calculated by DESeq2. Genes with log_2_fold change >1 and p_adjusted_-value <0.05 averaged across the two biological replicates (technical triplicate). These results indicate 57 genes significantly downregulated and 7 genes significantly upregulated. n=3 for each set of MEFs, technical replicate. Comparison is between aligned single-end reads from *Rtf2^+/+^* and *Rtf2^-/lox^* SV40-immortalized MEFs transduced with Hit & Run pMMP Cre retrovirus 72 hrs prior to harvest.^44^

**Extended Data Figure 5.**
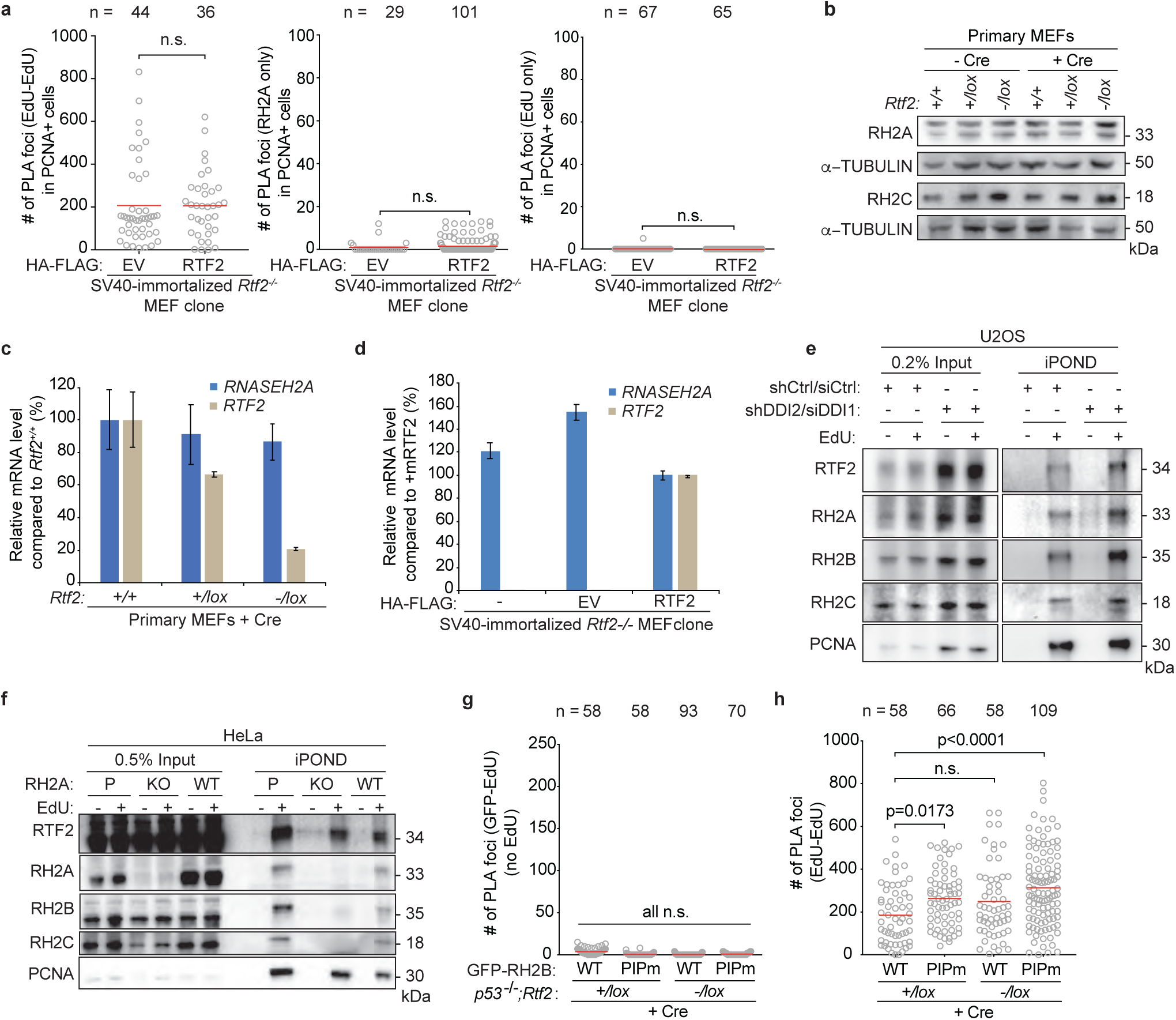
RTF2 deficiency results in loss of RNase H2 from the replication fork. **a,** Quantification of EdU-EdU PLA foci and single antibody (RNASEH2A and EdU) controls for nPLA in Fig. 2d. **b,** Representative immunoblot of whole cell lysates showing RNASEH2A and RNASEH2C levels in primary MEFs transduced with Hit & Run Cre recombinase retrovirus 72 hrs before harvest. **c,d,** RT-qPCR analysis of relative mouse *Rnaseh2a* and *Rtf2* transcript levels in primary MEFs transduced with Hit & Run pMMP Cre retrovirus 72 hrs before harvest or in SV40-LT immortalized RTF2-deficient sub-cloned MEF lines expressing HA-FLAG empty vector (EV) or mRTF2 (RTF2) cDNA constructs, respectively^44^. Expression is normalized to β*-actin*. **e,** Representative iPOND immunoblot in DDI-depleted U2OS cells. Proteins purified using streptavidin beads recognizing nascent DNA were detected by western blot. DDI depletion increases levels of RTF2 and RNase H2 on nascent chromatin. **f,** Representative iPOND immunoblot in CRISPR-edited RNASEH2A KO or WT HeLa cells. Proteins purified using streptavidin beads recognizing nascent DNA were detected by western blot. RNase H2 loss does not diminish recruitment of RTF2 to nascent chromatin. **g,** Quantification of GFP (RNASEH2B)-EdU PLA foci in untreated cells (no EdU) for nPLA in Fig. 2e. **h,** Quantification of EdU-EdU PLA foci for nPLA in Fig. 2e. Experiments were conducted at least three times in biological replicates with consistent results for a,b,e,f. Error bars represent standard deviation. Mean is indicated with a red line for a,g. Experiment conducted twice in biological replicates with consistent results for g. Significance evaluated by Kruskal-Wallis ANOVA with a Dunn’s post-test. RH2A = RNASEH2A, P = Parental, KO = CRISPR-mediated RH2A knockout, EV = empty vector, RH2B = RNASEH2B, RH2C = RNASEH2C.

**Extended Data Figure 6.**
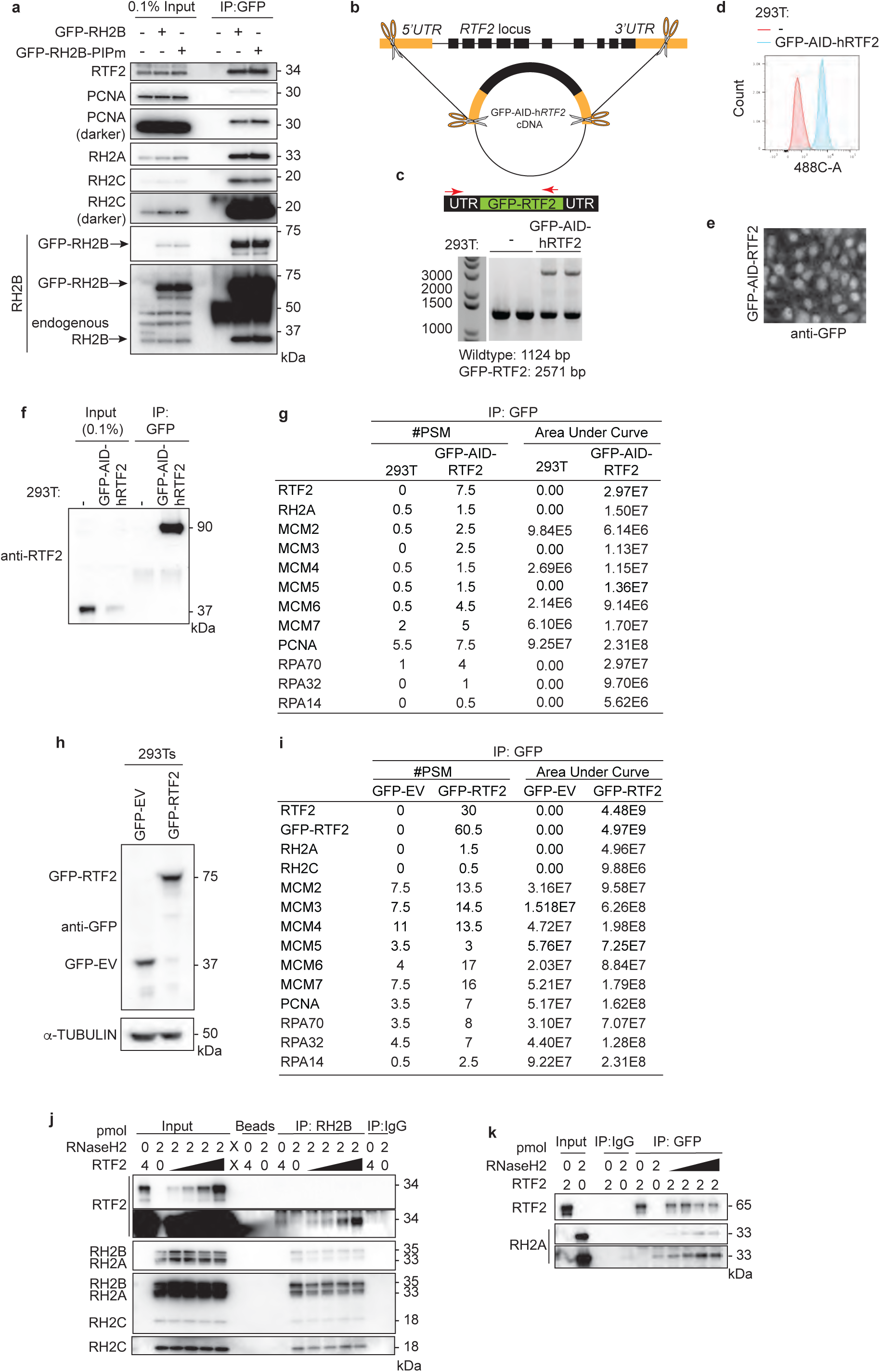
RTF2 interacts with RNase H2 and components of the replisome. **a,** Representative immunoblots following GFP immunoprecipitation of exogenously-expressed GFP-RNASEH2B-WT or GFP-RNASEH2B-PIP mutant from HEK293T cells. **b,** Schematic of CRISPR-Cas9 targeting to generate a tagged RTF2 construct expressed from the endogenous *RTF2* locus. A plasmid carrying a GFP-AID-h*RTF2* cDNA flanked by homology arms to the 5’UTR and 3’UTRs (orange boxes) of *RTF2* was targeted to the endogenous locus of Rtf2 in HEK293T cells and subsequently cloned. This line will be referred to as endogenous GFP-AID-RTF2 HEK293Ts. **c,** Genotyping analysis of wild type and endogenous GFP-AID-*RTF2* HEK293Ts. Schematic represents genotyping primers that were used to amplify the endogenously tagged locus. Forward primer recognizes *RTF2* promoter region upstream to the GFP-AID-h*RTF2* insert and reverse primer recognizes an *RTF2* exonic region. The primer pair amplifies a wild type *RTF2* allele of 1124 bp and the GFP-tagged allele of 2571 bp. **d,** Flow cytometry analysis of wild type and endogenous GFP-AID-*RTF2* HEK293Ts. **e,** Representative images of immunofluorescence analysis of endogenous GFP-AID-*RTF2* HEK293Ts. Cells were fixed and stained with anti-GFP antibodies. **f,** Representative immunoprecipitation with GFP antibodies from wild type and endogenous GFP-AID-*RTF2* HEK293Ts. Immunoblotted with RTF2 antibody. The GFP-AID-h*RTF2* protein is predicted to be 90.5 kDa. **g,** GFP was immunoprecipitated with GFP antibodies from wild type and endogenously targeted GFP-AID-RTF2 HEK293Ts in a-e. Average peptide spectral matches (#PSM) and area under the curve (AUC) from LC-MS for given proteins averaged across two biological replicates. **h,** Representative immunoprecipitation with GFP from HEK293Ts retrovirally expressing GFP-Empty Vector (GFP-EV) or GFP-human-RTF2 (GFP-RTF2). The GFP-hRTF2 construct is predicted to be 63.8 kDa. **i,** GFP was immunoprecipitated from the chromatin fraction of cell lysates with GFP nanobodies isolated from cells in g. Average #PSM and AUC from LC-MS for given proteins averaged across two biological replicates **j,** Representative immunoblot from immunoprecipitation of recombinant RNase H2 complex and RTF2 expressed in *E. coli*. Protein amount (pmol) are indicated above each lane; range of RTF2 is 0.5, 1, 2, 4 pmol. The blot corresponds to immunoblot shown in Figure 2f. **k,** Representative immunoblot from biochemical immunoprecipitation of recombinant GFP-tagged RTF2 and RNase H2 complex expressed in *E. coli*. Protein amount (pmol) indicated above each lane; range of RTF2 is 0.5, 1, 2, 4 pmol. Experiments were conducted at least two times in biological replicates with consistent results for e,g,i,j,k,l.

**Extended Data Figure 7.**
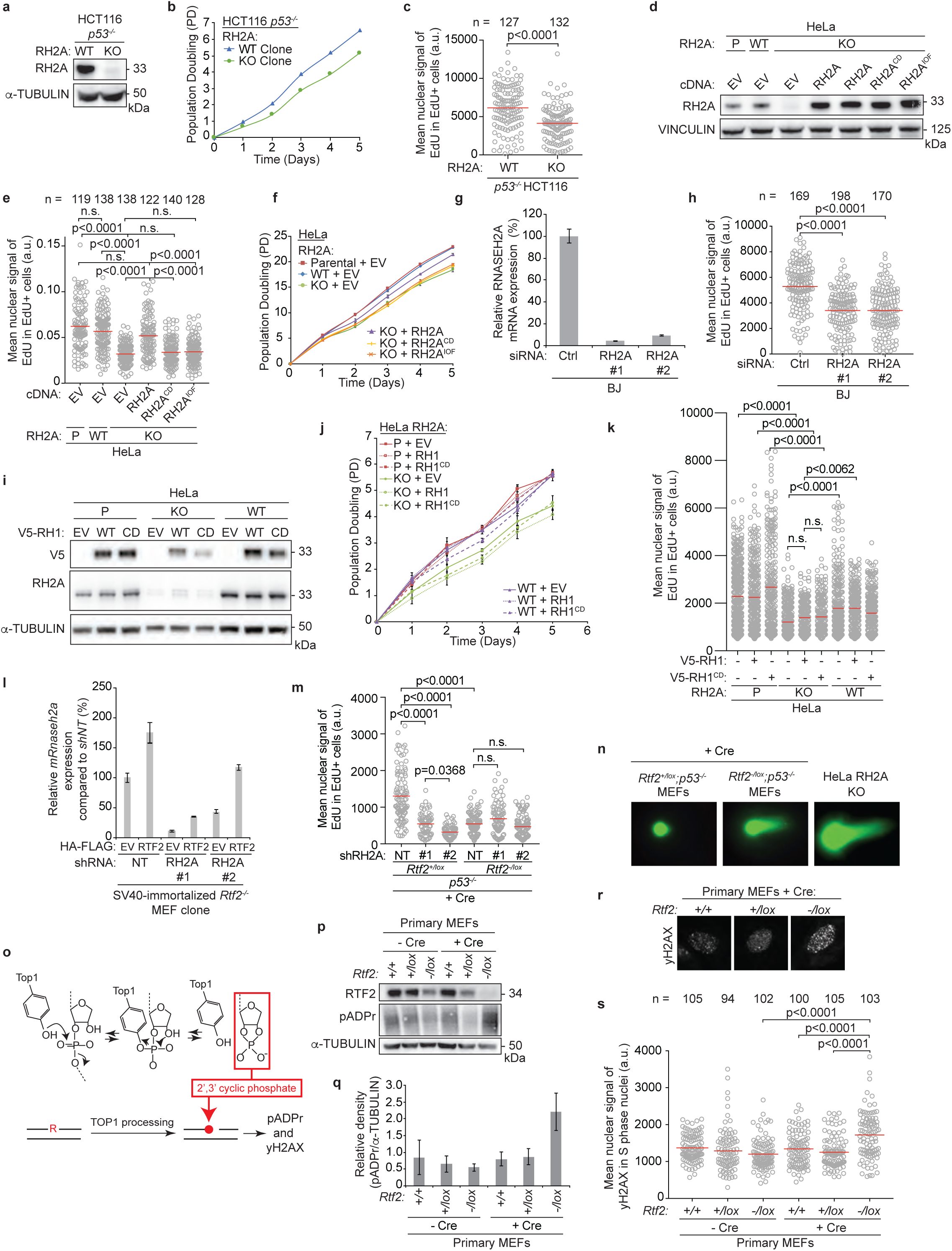
RNase H2-deficient cells phenocopy growth and replication phenotypes of RTF2 deficiency. **a,** Representative immunoblot showing loss of RNASEH2A in CRISPR-edited RNASEH2A KO HCT116 *p53^-/-^* cells. α-tubulin represents loading control. **b,** Growth curves of indicated cells. **c,** Quantification of representative experiment of mean signal of EdU in EdU-positive cells in indicated cells. **d,** Representative immunoblot showing complementation of wildtype, catalytic dead (RNASEH2A^D34A;D169A^), or isolation of function (RNASEH2A^P40D;Y210A^) RNASEH2A in CRISPR-edited RNASEH2A KO HeLa cells. Vinculin represents loading control. **e,** Quantification of representative experiment of mean signal of EdU in EdU-positive cells in indicated cells. **f,** Growth curves of indicated cells. **g,** Representative RT-qPCR analysis of relative human *RNASEH2A* transcript levels in BJ cells. Expression is normalized to *β-actin* expression. **h,** Mean nuclear signal of EdU in EdU-positive BJ cells treated with indicated siRNAs. **i,** Representative immunoblot showing expression of empty vector (EV), wildtype V5-RNASEH1, or catalytic dead V5-RNASEH1^D210N^ in CRISPR-edited RNASEH2A KO or HeLa cells. α-tubulin represents loading control. **j,** Growth curves of indicated cells. **k,** Quantification of representative experiment of mean signal of EdU in EdU-positive cells in indicated cells. **l,** Representative RT-qPCR analysis of mouse *Rnaseh2* expression in SV40-immortalized *Rtf2^-/-^* MEF clones expressing empty vector (EV) or HA-FLAG-mRTF2 (RTF2) transduced with indicated shRNAs. *Rnaseh2* expression is normalized to β*-actin*. **m,** Quantification of representative experiment of mean signal of EdU in EdU-positive cells in indicated cells. **n,** Representative images of neutral comet assay post RNase HII-digestion with olive tail moment quantified in Fig. 2i. **o,** Schematic of TOP1-mediated ribonucleotide processing in the absence of RNase H2. **p,** Representative immunoblot of whole cell lysates in primary MEFs 72 hrs after infection with Hit & Run Cre showing poly-ADP-ribosylation. **q,** Average densitometry plots from three biological replicates as in o. **r,** Representative immunofluorescent images of γH2AX staining in primary MEFs after infection with Hit & Run Cre. **s,** Quantification of representative experiment of mean nuclear signal of γH2AX from r. Experiments were conducted at least three times in biological replicates with consistent results for a,c,d,e,g,h,i,k,l,m,n,p,r,s. Experiment were conducted three times in biological replicates with technical triplicates for b,f,j. Error bars represent standard deviation. Each dot represents one cell for c,e,h,k,m,s. Mean for each sample shown with red line for c,e,h,k,m,s. Cells were pulsed with EdU for 1 hr prior to fixation for c,e,h,k,m,s. Experiments were blinded prior to analysis for c,h. Significance evaluated by Kruskal-Wallis ANOVA with a Dunn’s post-test. RH2A = RNASEH2A, P = Parental, WT = wildtype, EV = empty vector, RH2A^CD^ = catalytic dead RNASEH2A^D34A/D169A^, RH2A^IOF^ = isolation of function RNASEH2A^P40D/Y210A^, RH1 = RNASEH1, CD = catalytic dead RNASEH1^D210N^, Ctrl = Control, NT = Non-targeting.

**Extended Data Figure 8.**
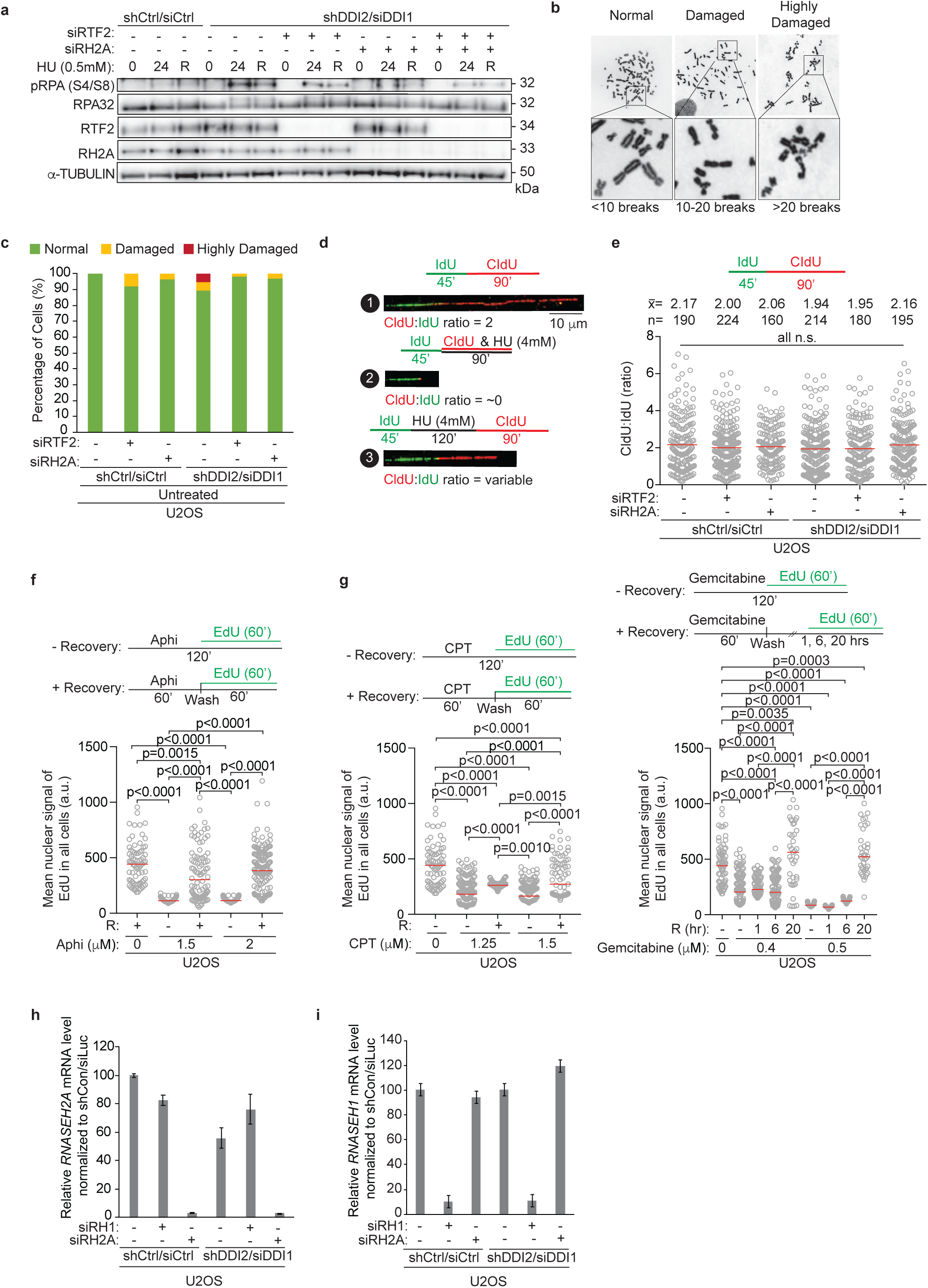
Removal of RNase H2, but not RNase H1, from stalled replication forks allows for replication restart and genome stability. **a,** Representative immunoblot in U2OS cells transduced or transfected with indicated RNAi reagents and treated with HU (0 = untreated, 24 = 24 hr treatment, R = 24 hr treatment followed by 8 hr release). **b,** Representative DNA metaphase spreads from U2OS cells after replication in the presence of low-dose aphidicolin for 40 hrs. Metaphase spreads were categorized into three classes of breakage severity: normal (<10 breaks), damaged (10-20 breaks), and highly damaged/uncountable (>20 breaks). **c,** Quantification of representative experiment of the percentage of metaphase spreads in each category described in **b** in U2OS cells transduced or transfected with indicated RNAi reagents (112 hrs). n>50 for metaphases scored in each sample. **d,** Schematic and representative images of DNA combing replication fork restart assay. **e,** Top: Labeling schematic. Bottom: Ratio of CldU tract to IdU tract lengths in U2OS cells infected or transfected with indicated siRNAs for 72 hrs. **f,** Top: Labeling schematic. Bottom: Mean nuclear signal of EdU in EdU-positive U2OS cells treated as indicated. **g,** Top: Labeling schematic. Mean nuclear signal of EdU in EdU-positive U2OS cells treated as indicated. **h,i,** RT-qPCR analysis of *RNASEH2A* or *RNASEH1* expression in indicated cells normalized to *GAPDH*. Experiments conducted at least three times in biological replicates with consistent results for a,b,c,e,f,g,h,i. Error bars represent standard deviation. Mean shown with red line for e,f,g. Experiments were blinded prior to analysis for g. Average CldU:IdU ratios are listed above each sample for g. Outliers removed with ROUT (1%) for g. Significance evaluated by Kruskal-Wallis ANOVA with a Dunn’s post-test. RH2A= RNASEH2A, Ctrl = Control. RH2A = RNASEH2A, Aphi = Aphidicolin, CPT = Camptothecin.

**Extended Data Figure 9.**
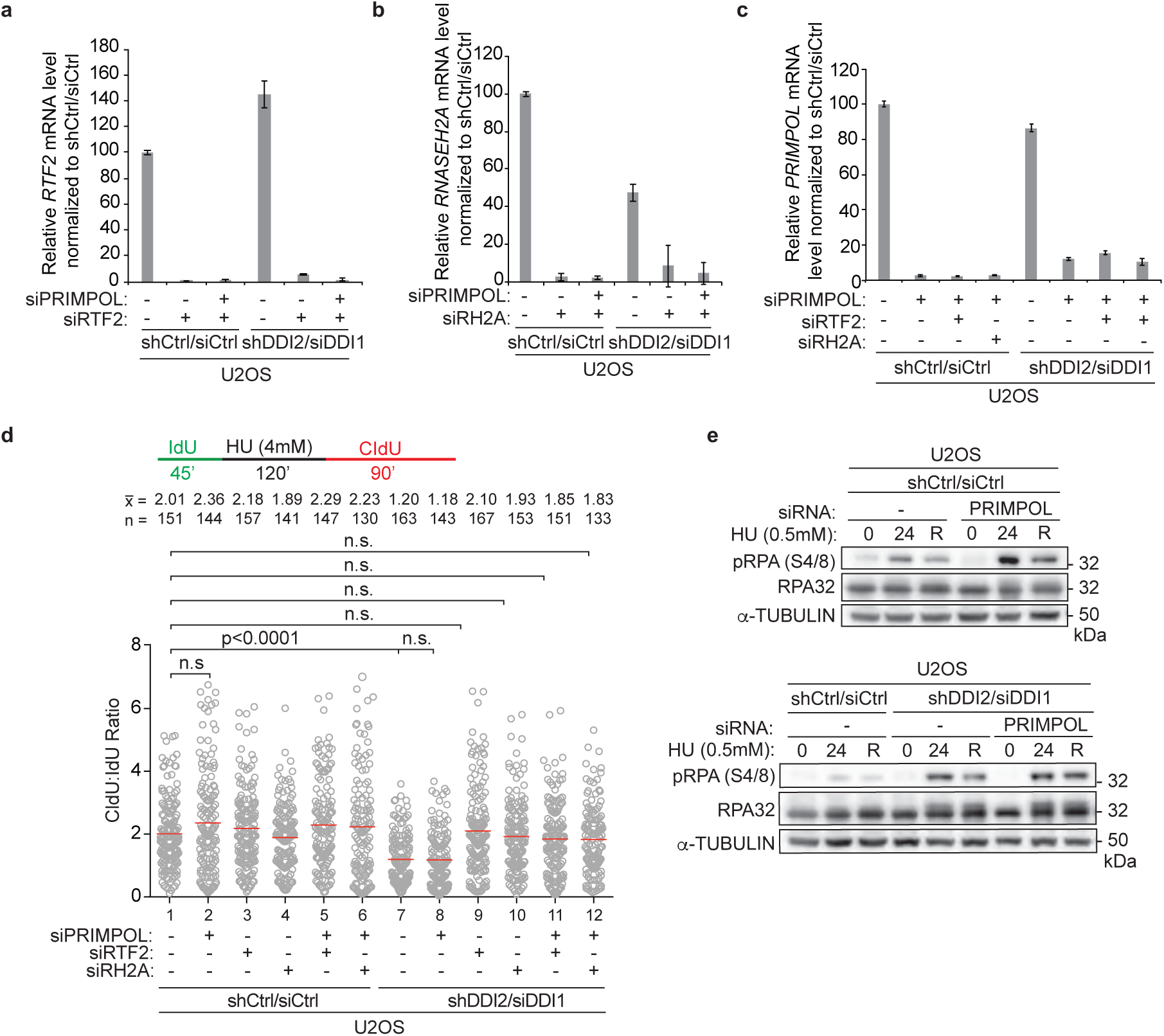
PRIMPOL deficiency induces p-RPA S4/8 signal, but not a restart efficiency defect in response to replication stress. **a-c,** Representative RT-qPCR analysis of relative human *RNASEH2A, RTF2,* or *PRIMPOL* transcript levels in indicated U2OS cells. Expression is normalized to *GAPDH*. **d,** Top: Labeling schematic. Bottom: Ratio of CldU:IdU tract lengths in indicated U2OS cells. **e,** Representative immunoblot in indicated U2OS cells treated with HU (0 = untreated, 24 = 24 hr treatment, R = 24 hr treatment followed by 8 hr release). α-tubulin represents loading control. Experiments were conducted at least three times in biological replicates with consistent results for a,b,c,d,e. Error bars represent standard deviation. Experiments were blinded prior to analysis for a,b,c,d,e. Mean is shown with red line for d. Average CldU:IdU ratios are listed above each sample for d. Outliers removed with ROUT (1%) for d. Significance evaluated by Kruskal-Wallis ANOVA with a Dunn’s post-test. RH2A= RNASEH2A, Ctrl = Control.

**Extended Data Figure 10.**
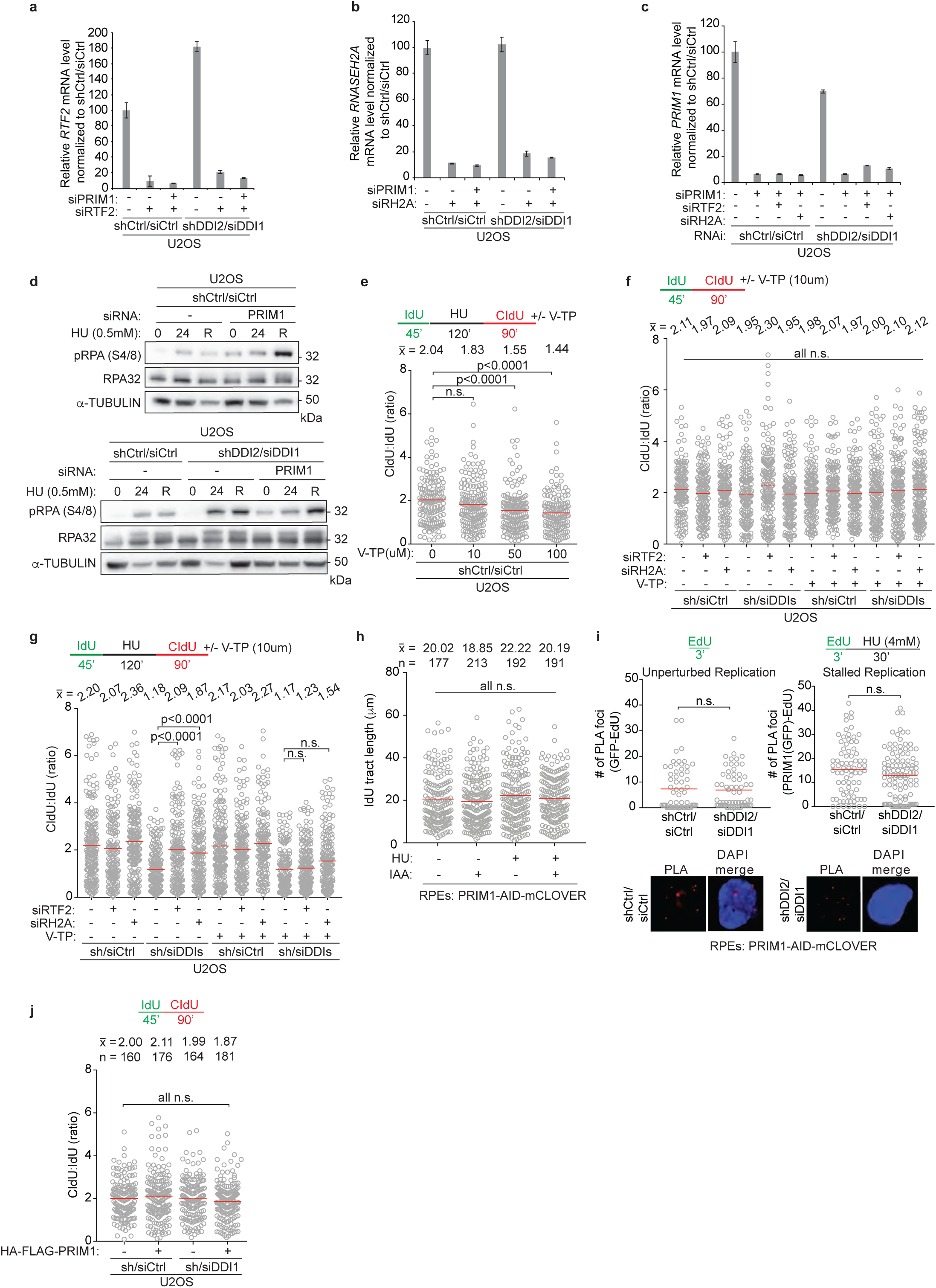
Inactivation or depletion of PRIM1 causes inefficient replication restart and induces p-RPA S4/8 signal. **a-c,** Representative RT-qPCR analysis of relative human *RNASEH2A, RTF2,* or *PRIM11* transcript levels in indicated U2OS cells. Expression is normalized to *GAPDH*. **d,** Representative immunoblot in U2OS cells transfected with siRNA and treated with HU as indicated (0 = untreated, 24 = 24 hr treatment, R = 24 hr treatment followed by 8 hr release). α-tubulin represents loading control. **e,f,g,** Top: Labeling schematics for DNA combing restart and progression assays in the setting of varying concentrations of V-TP, a potent PRIM1 inhibitor. Bottom: Quantification of representative experiment of CldU:IdU tract length ratios in U2OS cells transduced or transfected with indicated RNAi reagents. **h,** IdU lengths in indicated RPE cells from experiment in Fig. 4e. Mean for each sample shown with red line. Average length indicated above each sample. **i,** Top: nPLA labeling schematic. Middle: Quantification of PRIM1(GFP)-EdU PLA foci in PRIM1-AID-mCLOVER RPE cells in the setting of progressing or stalled replication. Bottom: Representative images of GFP(PRIM1)-EdU foci in PRIM1-AID-mCLOVER RPE cells. DAPI stains the nucleus. **j,** Top: Labeling schematic for DNA combing. Below: Quantification of representative experiment of CldU:IdU tract length ratios in U2OS cells transduced with either HA-FLAG-PRIM1 or HA-FLAG-EV and transduced or transfected with indicated RNAi reagents. Experiments were conducted at least three times in biological replicates with consistent results for a,b,c,d,e, f,g,h,i,j. Error bars represent standard deviation. Mean shown with red line for e,f,g,h,i,j. Experiments were blinded prior to analysis for e,f,g,h,j. Average CldU:IdU ratios or tract length are listed above each sample for e,f,g,h,j. Outliers removed with ROUT (1%) for e,f,g,h,j. Significance evaluated by Kruskal-Wallis ANOVA with a Dunn’s post-test. Ctrl = Control, RH2A = RNASEH2A, dox = doxycycline, V-TP = vidarabine.

