## Supplemental information for "RTF2 controls replication repriming and ribonucleotide excision at the replisome"

Supplementary Information includes:

| | |
|---|---|
| Supplementary Table 1. | Mouse strains |
| Supplementary Table 2. | Cell lines |
| Supplementary Table 3. | Primers |
| Supplementary Table 4. | siRNAs and shRNAs |
| Supplementary Table 5. | Plasmids |
| Supplementary Table 6. | Antibodies |

Supplementary table 1.  Mouse strains

| MOUSE STRAIN | SOURCE | IDENTIFIER |
|---|---|---|
| Rtf2 $^{tm1a(KOMP)wtsi}$ | This paper | |
| B6.Cg Tg(ACTFLPe)9205Dym/J | Jackson Laboratories | 005703 |
| B6.FVB/N-Tg(Ella-cre)C5379Lmgd/J | Jackson Laboratories | 003314 |
| C57BL/6J | Jackson Laboratories | 000664 |

Supplementary table 2. Mammalian cell lines

| **CELL LINE** | **SOURCE** | **IDENTIFIER** |
|---|---|---|
| Human: HEK 293T | ATCC | |
| Human: HEK 293T endogenously GFP-tagged RTF2 | This paper | N/A |
| Human: BJ-hTERT-E6/7, male | Smogorzewska Lab | |
| Human: U2OS, female | ATCC | HTB-96 |
| Human: RPE $p53^{-/-}$, $pRb^{-/-}$, PRIM1-AID-mClover | de Lange Lab | |
| Human: HeLa (HeLa Parental, HeLa WT Clone, HeLa RNASEH2A KO Clone), female | Durocher and Jackson Labs | |
| Human: HCT-116 $p53^{-/-}$ (HCT-116 $p53^{-/-}$ WT Clone, HCT-116 $p53^{-/-}$ RNASEH2A KO Clone) | Durocher and Jackson Labs | |
| Mouse: RTF2 MEFs ($Rtf2^{+/+}$, $Rtf2^{+/lox}$, $Rtf2^{-/lox}$, and $Rtf2^{lox/lox}$, SV40-immortalized $Rtf2^{-/-}$ clone, $p53^{-/-};Rtf2^{+/lox}$, $p53^{-/-};Rtf2^{-/lox}$) | This paper | N/A |
| KOMP mES cells $Rtf2^{tm1a(KOMP)wtsi}$ | KOMP | MGI code: 1913654 |

Supplementary table 3. Primers

| mESC long range genotyping primers | | |
|---|---|---|
| **Oligonucleotide** | **VENDOR** | **IDENTIFIER** |
| mESCs Cassette 3' Universal Forward cacacctccccctgaacctgaaac | IDT | KOMP |
| mESCs Cassette 5' Universal Reverse ggtggtgtgggaaagggttcgaag | IDT | KOMP |
| mESCs Cassette GF3 gccgaagaaggtcgagaaggtcag | IDT | KOMP |
| mESCs Cassette GR3 cgaatctctccacctgctcaatccag | IDT | KOMP |

| RTF2 mouse genotyping primers | | |
|---|---|---|
| **Oligonucleotide** | **VENDOR** | **IDENTIFIER** |
| Genotyping_PCR1_Fwd gcctgtgagcttggcaggta | IDT | This Paper |
| Genotyping_PCR1_Rev aggggaagacctgactgtgt | IDT | This Paper |
| Genotyping_PCR2_Fwd gcctgtgagcttggcaggta | IDT | This Paper |
| Genotyping_PCR2_Rev agcctgagctctgtcacatt | IDT | This Paper |
| Genotyping_PCR3_Fwd gatattgctgaagagcttgg | IDT | This Paper |
| Genotyping_PCR3_Rev gaagttatctcgacgaagttc | IDT | This Paper |

| RT-qPCR primers | | |
|---|---|---|
| **Oligonucleotide** | **VENDOR** | **IDENTIFIER** |
| Mouse *Rtf2* RT-qPCR Forward gaagtgtgtcacacgtgtgg | IDT | This Paper |
| Mouse *Rtf2* RT-qPCR Reverse tttcttttccagcttggccc | IDT | This Paper |
| Human *RTF2* RT-qPCR Forward tgctgaagacaaggatggag | IDT | Kottemann et al |
| Human *RTF2* RT-qPCR Reverse tgaaacagactctgctgcct | IDT | Kottemann et al. |
| Mouse *Rnaseh2a* RT-qPCR Forward gcatctttgccaaggtggcc | IDT | This Paper |
| Mouse *Rnaseh2a* RT-qPCR Reverse ggtcttgggatcattgggat | IDT | This Paper |
| Human *RNASEH2A* RT-qPCR Forward gctgaaagtggcagactcaa | IDT | This Paper |
| Human *RNASEH2A* RT-qPCR Reverse caggttgtatttgacccgcc | IDT | This Paper |
| Human *RNASEH1* RT-qPCR Forward aggaatcggcgtttactggg | IDT | This Paper |
| Human *RNASEH1* RT-qPCR Reverse aggctgcatgaatttccgc | IDT | This Paper |
| Human *DDI1* RT-qPCR Forward tggaacacaacgtgctacct | IDT | Kottemann et al. |
| Human *DDI1* RT-qPCR Reverse atctgtctgggggctgtct | IDT | Kottemann et al. |
| Human *DDI2* RT-qPCR Forward cgatgtagtgtgtgtgtactgc | IDT | Kottemann et al. |
| Human *DDI2* RT-qPCR Reverse ccagtgaggtagattctttaccactt | IDT | Kottemann et al. |
| Human *GAPDH* RT-qPCR Forward ggtcggagtcaacggattt | IDT | This Paper |
| Human *GAPDH* RT-qPCR Reverse gccccacttgattttggag | IDT | This Paper |
| Mouse b-actin RT-qPCR Forward ctaaggccaaccgtgaaaag | IDT | Thongthip et al. |
| Mouse b-actin RT-qPCR Reverse accagaggcatacagggaca | IDT | Thongthip et al. |
| Human *PRIM1* Forward RT-qPCR gacagagcattgaaggagga | IDT | This Paper |
| Human *PRIM1* Reverse RT-qPCR cgtcttgaccaccctttaca | IDT | This Paper |
| Human *PRIMPOL* Forward RT-qPCR ggcacttcagtagaaaccat | IDT | This Paper |

| | | |
|---|---|---|
| Human *PRIMPOL* Reverse RT-qPCR cgccgaattcctcctttaat | IDT | This Paper |

| Gateway cloning primers | | |
|---|---|---|
| **Oligonucleotide** | **VENDOR** | **IDENTIFIER** |
| attB human RNASEH2A Forward ggggacaagtttgtacaaaaaagcaggcttcatggatctcagcgagctgga | IDT | This Paper |
| attB human RNASEH2A Reverse ggggaccactttgtacaagaaagctgggtctagaggctggttgctgact | IDT | This Paper |
| attB mouse RTF2 Forward ggggacaagtttgtacaaaaaagcaggcttcatgggttgcgacggaggcac | IDT | This Paper |
| attB mouse RTF2 Reverse ggggaccactttgtacaagaaagctgggttcagaagcagtaggatgtgt | IDT | This Paper |
| attB human PRIM1 Forward ggggacaagtttgtacaaaaaagcaggcttcatggagacgtttgaccccac | IDT | This Paper |
| attB human PRIM1 Reverse ggggaccactttgtacaagaaagctgggtttatttctcaaggaaaattt | IDT | This Paper |

| Mutagenesis cloning primers | | |
|---|---|---|
| **Oligonucleotide** | **VENDOR** | **IDENTIFIER** |
| human RNASEH2B_F300A;F301A Forward ccaattttttttttattttttaccccagcagcggtatcaatacttttcattccactcttgtcaactttagcca | IDT | This Paper |
| human RNASEH2B_F300A;F301A Reverse tggctaaagttgacaagagtggaatgaaaagtattgataccgctgctggggtaaaaaataaaaaaaaaaattgg | IDT | This Paper |
| human RNASEH2A_D34A cctgggcgtcgctgaggcgggca | IDT | This Paper |
| human RNASEH2A_D169A caaggccaaagcagctgccctctacccgg | IDT | This Paper |
| human RNASEH2A_P40D cgggcaggggcgacgtgctgggcc | IDT | This Paper |
| human RNASEH2A_R210A actgattatggctcaggcgcccccaatgatcccaagac | IDT | This Paper |

| CRISPR cloning primers | | |
|---|---|---|
| **Oligonucleotide** | **VENDOR** | **IDENTIFIER** |
| RTF2 5'BamHI aaaaggatcccatgggttgcgacgggggga | IDT | This paper |
| RTF2 3'NotI aaaagcggccgctcagaagcagtaggacgtgtgg | IDT | This paper |
| hRTF 5’ UTR Fwd INFUSION accatgattacgccaagcttactcttggaacgggcatggc | IDT | This paper |
| hRTF 5’ UTR Rev INFUSION tcctcgcccttgctcaccatcgcaggacgggagtcagagc | IDT | This paper |

| | | |
|---|---|---|
| GFP Fwd INFUSION<br>atggtgagcaagggcgagga | IDT | This paper |
| hRTF rev INF<br>gagcggtggcagtgcgggcttcagaagcagtaggacgtgt | IDT | This paper |
| hRTF 3' UTR Fwd INF<br>agcccgcactgccaccgctc | IDT | This paper |
| hRTF 3' UTR Rev INF<br>aacgacggccagtgaattctaacttataggcagataaaat | IDT | This paper |
| mouse sgTrp53 exon 5 Fwd<br>caccgaagtcacagcacatgacgg | IDT | This paper |
| mouse sgTrp53 exon 5 Rev<br>aaacccgtcatgtgctgtgacttc | IDT | This paper |

Supplementary table 4. siRNAs and shRNAs

| **Oligonucleotide** | **VENDOR** | **IDENTIFIER** |
|---|---|---|
| Luciferase siRNA (siCtrl) | Thermo Fischer | 12935146 |
| hRNASEH1 siRNA 1 gggaaagaggugaucaacatt | Ambion | s48356 |
| hRNASEH1 siRNA 2 cagacaguauguuuacgautt | Ambion | s48357 |
| hRNASEH1 siRNA 3 cgggauuuauaggcaauatt | Ambion | s48358 |
| hRNASEH2A siRNA 1 caaugaucccaagacaaatt | Ambion | s20656 |
| hRNASEH2A siRNA 2 ccaccgauauuuccuggaatt | Ambion | s20657 |
| hRTF2 siRNA 1 caaagaugccgucauugaatt | Ambion | s226737 |
| hPrim1 siRNA 1 gaaccagagauuauaagaatt | Ambion | s11050 |
| hPrim1 siRNA 2 caucgucucuggguauauutt | Ambion | s11051 |
| hPrim1 siRNA 3 caacuacgguggagugauatt | Ambion | s10052 |
| hPrimpol siRNA 1 ggcuauugauagaguuaaatt | Ambion | s11053 |
| hPrimpol siRNA 2 ccacgaagaagagaucauatt | Ambion | s11054 |
| hPrimpol siRNA 3 ggaucuccuucgauuuagatt | Ambion | s11055 |
| hDDI1 siRNA1 ccggagacaucaauguuccaucgat | ThermoFisher | HSS181016 |
| hDDI1 siRNA 2 ggaaauuacacauucagucauggat | ThermoFisher | HSS140552 |
| hDDI1 siRNA 3 ccggagacaucaauguuccaucgat | ThermoFisher | HSS140553 |
| hDDI2 shRNA uggaaaucgauacagcuca | Open Biosystems | V3LHS_328065 |
| mRNASEH2A shRNA #1 ccgggctcgattacaacagcactttctcgagaaagtgctgttgtaatcgagctttttg | MilliporeSigma | TRC0000119585 |
| mRNASEH2A shRNA #2 ccggcgggtcgttgtcgtctgagttctcgagaactcagacgacaacgacccgtttttg | MilliporeSigma | TRC0000119584 |

Supplementary Table 5. Plasmids

| Plasmid | VENDOR | IDENTIFIER |
|---|---|---|
| ***Gateway Entry Vectors*** | | |
| pDONOR233; spectinomycin resistant; Gateway | Hill and Vidal Labs | pDONOR233 |
| pENTR223-EV; spectinomycin resistant; Gateway | Smogorzewska Lab | AS183 |
| pENTR223-mRTF2; spectinomycin resistant; Gateway | This paper | BC153 |
| pENTR223-RNASEH2A; spectinomycin resistant; Gateway | This paper | NJB2 |
| pENTR223-RNASEH2A$^{CD}$_D34A;D169A; spectinomycin; Gateway | This paper | NJB6 |
| pENTR223-RNASEH2A$^{IOF}$_P40D;Y210A; spectinomycin; Gateway | This paper | NJB11 |
| pENTR223-RNASEH2B; spectinomycin resistant; Gateway | This paper | NJB27 |
| pENTR223-RNASEH2B$^{PIPm}$_ F300A;F301A; spectinomycin resistant; Gateway | This paper | NJB29 |
| pENTR223-hPRIM1; spectinomycin resistant; Gateway | This paper | CB35 |
| ***Retroviral expression vectors*** | | |
| PEA59-GFP-EV-dest (destination vector); chloramphenicol and ampicillin resistant; Gateway | Smogorzewska Lab | AS769 |
| PEA59-GFP-EV-puro; ampicillin resistant; retroviral; Gateway | Smogorzewska Lab | AS1050 |
| PEA59-GFP-mRTF2-puro; retroviral; ampicillin resistant; Gateway | This paper | BC123 |
| pMSCVpuro-DEST (destination vector); chloramphenicol and ampicillin resistant; Gateway | Addgene | Plasmid# 119745 |
| pMSCVpuro-EV; ampicillin; Gateway | This paper | NJB22 |
| pMSCVpuro-RNASEH2A; ampicillin; Gateway | This paper | NJB18 |
| pMSCVpuro-RNASEH2A-siRNA Resistant; ampicillin; Gateway | This paper | NJB19 |
| pMSCVpuro-RNASEH2A$^{CD}$_D34A;D169A -siRNA Resistant; ampicillin; Gateway | This paper | NJB20 |
| pMSCVpuro- RNASEH2A$^{IOF}$_P40D;Y210A-siRNA Resistant; ampicillin; Gateway | This paper | NJB20 |
| pMSCV_PM_shRNA_Control_puro | Elledge lab | BC75 |
| pMSCV_PM_shRNA_shDDI2_puro | Smogorzewska Lab | MK59-63 |
| pMSCV-GFP-H2B-hygro | Smogorzewska Lab | YK77 |
| pEGFP-RNASEH2B; kanamycin resistant; Gateway | Addgene | Plasmid #108697 |
| pMMP Hit & Run Cre; retroviral; self-excising | Livingston Lab | NA |
| pWZL Cre-hygro; retroviral | de Lange Lab | NA |

| | | |
|---|---|---|
| pMSCV-HA-FLAG-Dest: destination vector; chloramphenicol and ampicillin resistant; Gateway | Smogorzewska Lab | AMS157 |
| pMSCV-HA-FLAG-EV: ampicillin resistant; Gateway | Smogorzewska Lab | AMS184 |
| pMSCV-HA-FLAG-hPRIM1: ampicillin resistant; Gateway | This paper | CB36 |
| ***Lentiviral expression vectors*** | | |
| pLKO.1 shRNASEH2A #1_puro; ampicillin resistant | MilliporeSigma | SHCLNG-NM_027187, TRC0000119585 |
| pLKO.1 shRNASEH2A #2_puro; ampicillin resistant | MilliporeSigma | SHCLNG-NM_027187, TRC0000119584 |
| pLKO.1 shRNA Control Plasmid puro; ampicillin resistant; | MilliporeSigma | SHC002 |
| pLVpuro-CMV-N-EGFP; destination vector; chloramphenicol and ampicillin resistant; Gateway | Addgene | Plasmid #122848 |
| pLVpuro-CMV-N-EGFP- RNASEH2B; ampicillin resistant; Gateway | This paper | PR3 |
| pLVpuro-CMV-N-EGFP-RNASEH2B$^{PIPm}$_F300A;F301A; ampicillin resistant; Gateway | This paper | PR5 |
| ***Other expression vectors*** | | |
| ppyCAG_RNaseH1_WT; ampicillin resistant | Addgene | Plasmid #111906 |
| ppyCAG_RNaseH1_D210N; ampicillin resistant | Addgene | Plasmid #11904 |
| ***CRISPR generation of GFP-AID-RTF2 HEK 293Ts*** | | |
| pcDNA5-FRT-TO-EGFP-AID | Addgene | Plasmid #80075 |
| MLM3636 | Addgene | Plasmid #43860 |
| MLM3636 5' UTR_1 sequence acgctaggcgcggcgtagcg | This paper | BC201, 202 |
| px330 | Addgene | Plasmid #42230 |
| px330 3'UTR_1 atgtgaggcgtgtcggttcc | This paper | BC204, 206 |
| pUC19-5UTR-GFP-AID-hRTF2-3UTR Homology Donor Construct | This paper | BC219 |
| ***CRISPR generation of p53-/-;Rtf2$^{+/lox}$ and p53-/-;Rtf2$^{-/lox}$ MEFs*** | | |
| pSpCas9(BB)-2A-Puro (PX459) V2.0 | Addgene | Plasmid #62988 |
| pX459-sgTRP53 | This paper | NJB92 |
| ***Protein Biochemistry*** | | |
| PSKA002 HIS14-SUMO-MCS Expression Vector | Klinge Lab | |
| PSKA002 HIS14-SUMO-RTF2 | This paper | BC142 |

| | | |
|---|---|---|
| PSKA008 HIS14-GFP-MCS-Expression Vector | Klinge Lab | |
| PSKA008 HIS14-GFP-RTF2 | This paper | BC138 |
| pGEX6P1-hsRNASEH2BCA | Addgene | Plasmid #108692 |
| ***Protein Biochemistry*** | | |
| VSV-G; retroviral packaging | Mulligan lab | AMS166 |
| Gagpol; retroviral packaging | Mulligan lab | AMS167 |
| pMD2.G (VSV-G envelope expressing plasmid); lentiviral packaging | Addgene | Plasmid #12259 |
| psPAX2; lentiviral packaging | Addgene | Plasmid #12260 |

Supplementary Table 6. Antibodies

| ANTIBODY | SOURCE | IDENTIFIER | LOT (if known) |
|---|---|---|---|
| ***Primary Antibodies*** | | | |
| Mouse IgG | Santa Cruz | Cat# sc-2025, RRID:AB_737182 | D2022 |
| Rabbit IgG | Santa Cruz | Cat# sc-2027, RRID:AB_737197 | |
| Rabbit IgG | Cell Signaling Technology | Cat# 2729, RRID:AB_1031062 | 10 |
| Mouse monoclonal anti-α-tubulin (clone DM1A), WB:1:5000 | MilliporeSigma | Cat# T9026, RRID:AB_477593 | 0000137585 |
| Mouse monoclonal anti-γH2AX Ser139 (clone JBW301), IF 1:2000 | MilliporeSigma | Cat# 05-636, RRID:AB_309864 | 3782118 |
| Mouse monoclonal anti-biotin, IF: 1:2000 | Jackson ImmunoResearch | Cat# 200-002-211, RRID:AB_2339006 | 151728 |
| Rabbit monoclonal anti-biotin, IF: 1:2000 | Bethyl | A150-109A | 11 |
| Mouse monoclonal anti-BrdU (B44), combing: 1:10 | BD Biosciences | Cat# 347580, RRID:AB_400326 | 9172603 |
| Mouse monoclonal anti-Poly (ADP-Ribose) Polymer antibody [10H], WB:1:100 | Abcam | Cat # ab14459, RRID:AB_301239 | |
| Mouse monoclonal anti-PCNA (PC10), WB: 1:1000 | Santa Cruz | Cat# sc-56, RRID:AB_628110 | K1121 |
| Mouse Monoclonal anti-vinculin, Unconjugated, Clone hVIN-1 | MilliporeSigma | Cat# V9131, RRID:AB_477629 | 018M4779V |
| Rat monoclonal anti-BrdU [BU1/75 (ICR1)], combing: 1:20 | Abcam | Cat# ab6326, RRID:AB_305426 | |
| Rabbit monoclonal anti-MCM7 (D10A11) XP, WB: 1:1000 | Cell Signaling Technology | Cat# 3735S, RRID:AB_2142705 | 3 |
| Rabbit polyclonal anti-c20orf43 (RTF2), WB: 1:500 | Novus | Cat# NBP2-30645 | R72074 |
| Rabbit polyclonal anti-RTF2, WB: 1:500 | Proteintech | Cat# 16633-1-AP, RRID: AB_2256547 | 00074586 |

| | | | |
|---|---|---|---|
| Mouse monoclonal anti-RTF2 (clone OTI1E8), WB: 1:1000 | LS Bio | Cat# LS-C340588 | 75213 |
| Rabbit polyclonal anti-GFP | Smogorzewska Lab | Kotteman et al. | |
| Rabbit polyclonal anti-GFP | Abcam | Cat# ab290 RRIDL AB_303395 | GR3251545-1 |
| Mouse monoclonal Anti-GFP, WB: 1:3000 | Roche | Cat# 11814460001 | |
| Rabbit polyclonal anti-RNASEH2A, WB: 1:500 | Proteintech | Cat# 16132-1-AP, RRID:AB_2269729 | 00023264 |
| Rabbit polyclonal anti-RNASEH2A, WB: 1:500 | Abcam | Cat# ab83943, RRID:AB_1861175 | GR3212381-12 |
| Mouse monoclonal anti-RNASEH2A, WB:1:500 | Santa Cruz | Cat# sc-515475 | |
| Rabbit polyclonal anti-RNASEH2C, WB: 1:500 | AbClonal | Cat# A13884, RRID:AB_2760737 | 0067820201 |
| Rabbit polyclonal anti-RNASEH2C, WB: 1:500 | Abcam | Cat# ab89726, RRID:AB_2042815 | GR74866-1 |
| Rabbit polyclonal anti-RNASEH2B, WB: 1:1000 | Thermo Fisher Scientific | Cat# PA5-59059, RRID:AB_2646610 | V13088599 |
| Rabbit polyclonal anti-RNASEH2B, WB: 1:1000 | Atlas Antibodies | Cat# HPA041469, RRID:AB_2677496 | 000043725 |
| Rabbit polyclonal anti-phospho-RPA32 (S4/8), WB:1:1000 | Bethyl | Cat# A300-245A, RRID:AB_210547 | 7 |
| Rabbit polyclonal anti-RPA32, WB 1:2000 | Bethyl | Cat# A300-244A, RRID:AB_185548 | 3 |
| Rabbit polyclonal anti-PRIM1, WB:1000 | Proteintech | Cat # 10773-1-AP RRID:AB_2237549 | 00009142 |
| Mouse monoclonal anti-HA.11 Epitope Tag, WB: 1:3000 | BioLegend | Cat# 901514, RRID:AB_2565336 | B272772 |

| | | | |
|---|---|---|---|
| Mouse monoclonal anti-V5 tag antibody [SV5-PK1], WB: 1:5000 | Abcam | (Abcam Cat# ab27671, RRID:AB_471093) | GR3337308-16 |
| Sheep polyclonal anti-human RNase H2 complex, WB: 1:500 | Jackson Lab | | |
| ***Secondary Antibodies*** | | | |
| Goat Anti-Mouse IgG H&L Cross-Absorbed (Alexa Fluor® 488), IF:1:1000, combing:1:100 | ThermoFisher | Cat# A-11001, RRID:AB_2534069 | 632115 |
| Goat Anti-Mouse IgG H&L Cross-Absorbed (Alexa Fluor® 647), combing:1:100 | ThermoFisher | Cat# A-21235, RRID:AB_2535804 | 1837146 |
| Goat Anti-Rabbit IgG H&L (Alexa Fluor® 488) | ThermoFisher | Cat# A-11008, RRID:AB_143165 | 645151 |
| Goat Anti-Rat IgG H&L Cross-Absorbed (Alexa Fluor® 594), IF: 1:1000, combing: 1:100 | ThermoFisher | Cat# A-11007, RRID:AB_10561522 | 2107787 |
| Peroxidase-AffiniPure Goat Anti-Mouse IgG (H + L) antibody, WB: 1:5000 | Jackson ImmunoResearch Labs | Cat# 115-035-003, RRID:AB_10015289 | |
| Peroxidase-AffiniPure Donkey Anti-Sheep IgG (H + L) antibody, WB: 1:5000 | Jackson ImmunoResearch Labs | Cat# 713-035-003, RRID:AB_2340709 | |
| Peroxidase-AffiniPure Goat Anti-Rabbit IgG (H + L) antibody, WB: 1:5000 | Jackson ImmunoResearch Labs | Cat# 115-035-144, RRID:AB_2307391 | |
